# Aging directs the differential evolution of KRAS-driven lung adenocarcinoma

**DOI:** 10.1101/2025.01.20.633951

**Authors:** Felicia Lazure, Stanislav Drapela, Xiaoxian Liu, John H. Lockhart, Hossein Kashfi, Nadir Sarigul, Didem Ilter, Elsa R. Flores, Xuefeng Wang, Inna Smalley, Alex Jaeger, Xiaoqing Yu, Ana P. Gomes

## Abstract

Lung adenocarcinoma (LUAD), the most common histological subtype of lung cancer(*1, 2*), is a disease of the elderly, with an average age of diagnosis of about 70 years of age(*3*). Older age is associated with an increased incidence of KRAS-driven LUAD(*4*), a particularly deadly type of LUAD characterized by treatment resistance and relapse. Despite this, our understanding of how old age shapes KRAS-driven LUAD evolution remains incomplete. While the age-related increase in cancer risk was previously ascribed to the accumulation of mutations over time, we are now beginning to consider the role of host biology as an independent factor influencing cancer. Here, we use single-cell RNA-Sequencing of KP (Kras^G12D/+^; Trp53^flox/flox^) LUAD transplanted into young and old mice to define how old age affects LUAD evolution and map the changes that old age imposes onto LUAD’s microenvironment. Our data demonstrates that the aged lung environment steers LUAD evolution towards a primitive stem-like state that is associated with poor prognosis. We ascribe this differential evolution, at least in part, to a population of rare and highly secretory damage-associated alveolar differentiation intermediate (ADI) cells that accumulate in the aged tumor microenvironment (TME) and that dominate the niche signaling received by LUAD cells. Overall, our data puts aging center stage in coordinating LUAD evolution, highlighting the need to model LUAD in its most common context and creating a framework to tailor future cancer therapeutic strategies to the age of the patient to improve outcomes in the largest and most vulnerable LUAD patient population, the elderly.

## Main

The aging process results in widespread changes in the body at the molecular, cellular, tissue and systemic levels(*5*). Given that cancer progression is guided by the selective pressures that the tumor’s environment exerts(*6*) we first sought to characterize the consequences of old age in the lung. Gene Set Enrichment Analysis (GSEA) of bulk RNA-Sequencing (RNA-Seq) data comparing young and old mouse lungs(*7*) revealed distinct transcriptional profiles in aged lungs, with enrichment for pathways related to UV response and p53 signaling (Fig. 1A, table S1) indicative of the accumulation of damage that occurs as a function of age, and is in line with the increase in senescence (Fig. 1B) and DNA damage (Fig. 1C) observed in the aged lungs. GSEA also revealed an enrichment in pathways related to immune function and inflammation, including NF-κB signaling (Fig. 1A, table S1), which aligns with an age-related imbalance in the proportions of immune cells, including CD4 T-cells, CD8 T-cells and NK cells (Fig. 1D and fig. S1A) in accordance with previous literature(*8, 9*) as well as with the age-induced increase in abundance of various cytokines present in bronchioalveolar lavage fluid (Fig. 1E). While epithelial and stromal cell proportions remain largely unaltered by old age (Fig. 1F and G, fig. S1B), aged lungs displayed an increase in collagen abundance (Fig. 1H), which highlights that the function of the aged stromal cells is likely changed and is consistent with the age-related increase in respiratory diseases such as idiopathic pulmonary fibrosis(*10*). Together, this data combined with literature(*8, 11, 12*) establishes the aged lung environment as a markedly different milieu compared to a young lung and raises the question of whether these changes alter the evolution of LUAD (Fig. 1).

**Figure 1.**
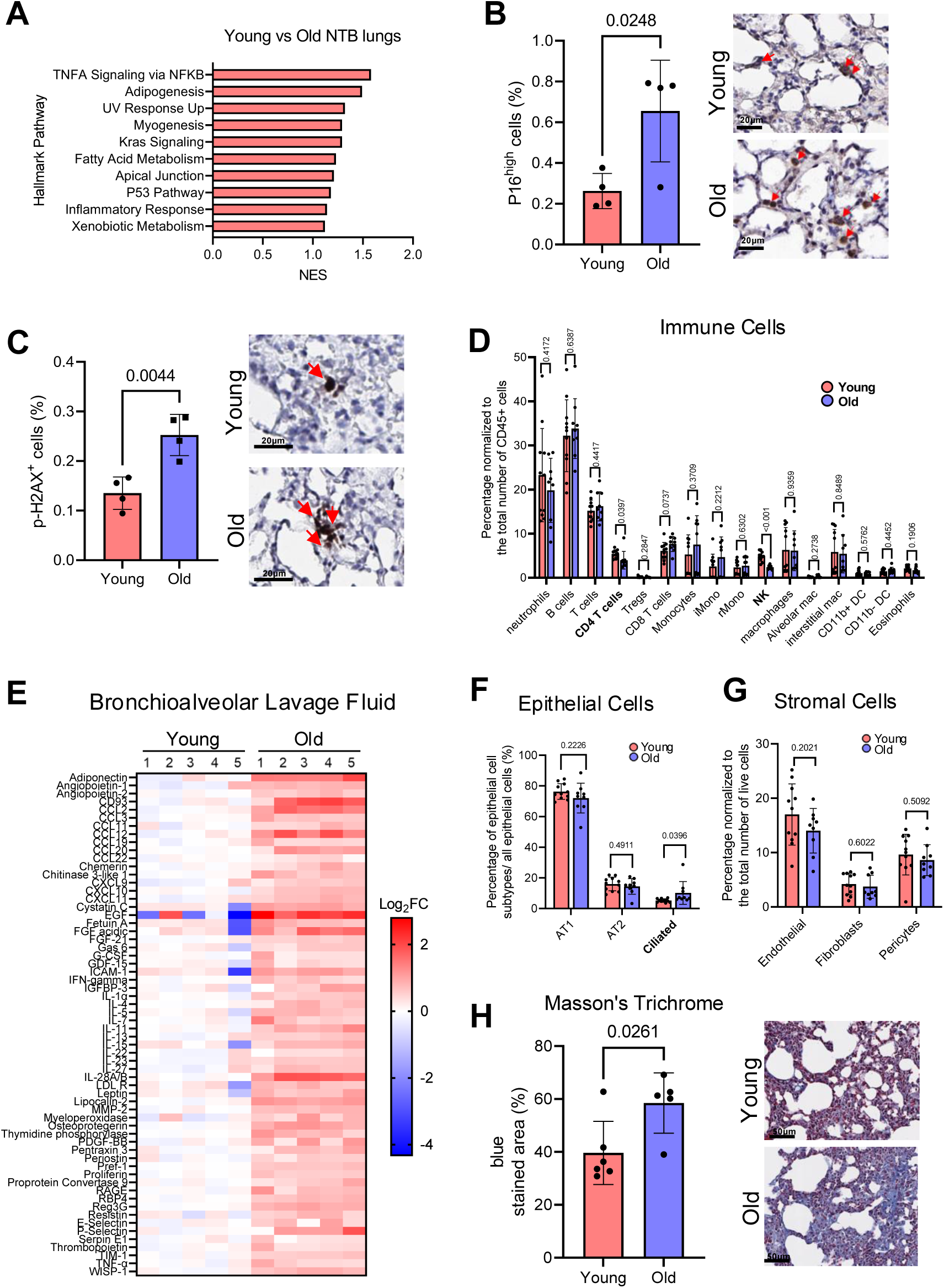
Old age alters the lung tissue landscape. **A.** Top 10 Hallmark pathways enriched in old (24-months) compared to young (6-months) non-tumor bearing lung bulk RNA-Seq samples(*7*) via Gene Set Enrichment Analysis (n=3 mice per group) **B.** Quantification of the percentage of P16^High^ cells in young (5-6 month) and old (22-24 month) non-tumor-bearing mouse lungs, along with representative IHC images (n=4 mice per group, t-test). Arrows represent P16^HIGH^ cells. **C.** Quantification of the percentage of p-H2A.x cells in young (5-6 month) and old (22-24 month) non-tumor-bearing mouse lungs along with representative IHC images (n=4 mice per group, t-test) Arrows represent p-H2AX^+^ cells. **D.** Flow cytometry immune cell type analysis in non-tumor-bearing lungs from young and old NTB mice (n=9-11 mice per group, t-test) **E.** Heat map of Log_2_FC of intensity values from a proteome profiler cytokine array ran on bronchioalveolar lavage fluid from young and old mice. (n=5 per group, t-test). **F.** Flow cytometry analysis of epithelial cell types in non-tumor-bearing lungs from young and old NTB mice. (n=9-11 mice per group, t-test) **G.** Flow cytometry analysis of stromal cell types in non-tumor-bearing lungs from young and old NTB mice. (n=9-11 mice per group, t-test) **H.** Quantification of blue area (ImageJ threshold hue:102-175, threshold saturation: 0-255, threshold brightness: 0-229) representing collagen and representative images of Masson’s trichrome staining of young (5-6 month) and old (22-24 month) non-tumor-bearing lungs. (n=5-6 mice per group, t-test) Data are presented as the mean ± SD.

To uncouple the involvement of age-driven cell-intrinsic changes from cell-extrinsic environmental selective pressures in LUAD, we optimized a syngeneic transplantation model (Fig. 2A and fig. S2). We injected lung cancer cells derived from the KP (Kras^G12D/+^; Trp53^flox/flox^)(*13*) genetically engineered mouse model (GEMM) of LUAD into WT C57BL/6 mice and allowed tumors to form in the lungs for 4 weeks (Fig. 2A). This model generated tumors with morphological(*14*) (fig. S2A) and transcriptional (fig. S2B and C) features similar to those of the KP GEMM(*15*), validating this transplantation system as an appropriate tool to study tumor evolution in different host mice. Moreover, unlike the autochthonous KP model, which upon induction promotes the recombination of pro-tumorigenic alleles in all AT2 cells(*13*), this transplantation model enables the interrogation of intratumoral and adjacent normal AT2 cells, which are a feature of human LUAD(*16*) and whose function significantly changes with aging(*12, 17*). Transplantation of KP cells into the lungs of young (5 – 6 months) and aged (19 – 24 months) mice did not produce significant differences in tumor burden, number or grade(*14*) between age groups (fig. S2D to H). Given that advanced age is linked to reduced LUAD initiation(*18*), these findings suggest that while the aged microenvironment may be less permissive to *de novo* tumorigenesis, it does not impede the growth of already transformed cells (fig. S2D to H). This raises the intriguing possibility that tumor initiation occurs significantly prior to diagnosis, and the trajectory and evolution of LUAD is shaped by the aged microenvironment well before diagnosis.

**Figure 2.**
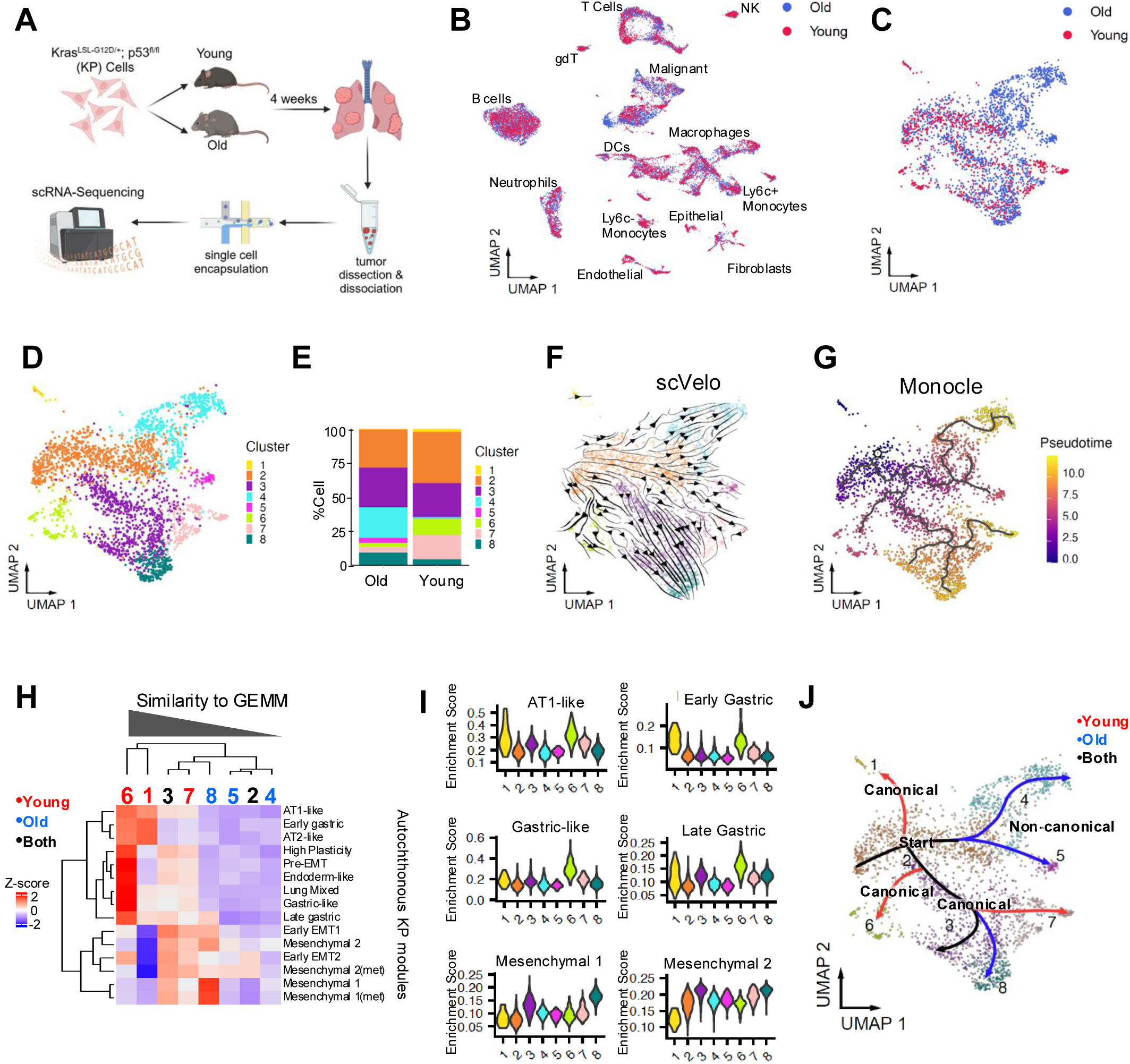
Aging drives the divergence of KRAS-driven LUAD evolution. **A.** Schematic of KP transplantation and scRNA-Seq experiment **B.** UMAP projection of all TME cells from micro dissected KP tumors from young and old mice, colored by age (n=3 mice per group) **C.** UMAP projection of KP epithelial malignant cells from young and old mice, colored by age (n=3 mice per group) **D.** UMAP projection of KP epithelial malignant cells from young and old mice, colored by cluster (n=3 mice per group) **E.** Bar graph of malignant epithelial cell cluster proportions in young and old mice (n=3 mice per group) **F.** scVelo trajectory analysis on the UMAP projection of epithelial malignant cells isolated from young and old mice, colored by cluster. Arrows represent the direction of evolution. (n=3 mice per group) **G.** scVelo trajectory analysis on the UMAP projection of epithelial malignant cells isolated from young and old mice, colored by pseudotime. (n=3 mice per group) **H.** heatmap of z-scores comparing our scRNA-Seq clusters of KP malignant cells isolated from young and old mice to the modules of KP evolution from Yang D., et al. *Cell* (2022) **I.** Violin Plots showing the enrichment score for select KP modules in clusters from young and old KP tumor-bearing mice **J.** UMAP of epithelial malignant cells with consensus arrows depicting evolutional trajectory in F and G and labeled based on matching with canonical KP modules in H and I.

To understand if aging alters the molecular landscape or evolution of LUAD, we next performed single-cell RNA-Sequencing (scRNA-Seq) on KP tumors micro-dissected from young and old mice (Fig. 2A and fig. S3). 30,547 cells were sequenced and all the expected major cell types(*8*) were recovered by our scRNA-seq analysis (Fig. 2B and fig. S3A). Notably, among all major cell types identified within the TME, the epithelial cell compartment, which contains the malignant KP cells transplanted, showed the most striking age-based segregation according to Unsupervised Uniform Manifold Approximation and Projection (UMAP) based dimensional reduction (Fig. 2B and C). Subsequent Louvain clustering further confirmed age-related shifts in malignant cell cluster proportions, unveiling common and unique subpopulations across ages (Fig. 2D and E, table S2). Trajectory inference analyses using scVelo(*19*) and Monocle(*20*) unveiled distinct trajectories of KP evolution, all originating from the common cluster 2 (Fig. 2F and G). This cluster also showed the highest correlation with cultured parental KP cells, reinforcing its role as the evolutionary starting point (fig. S4A). This trajectory analysis revealed two main paths of KP evolution that segregate by age. On one hand, we observe the canonical path of LUAD evolution previously established in the KP autochthonous model(*15*), where KP cells transitioned from cluster 2 to cluster 3, which then gives rise to terminal branches 6, 7 and 8 whose predominance depends on the age of the lung TME (Fig. 2D to G). To define the properties of each cluster, we compared them to known modules of canonical KP evolution in the autochthonous model(*21*) (Fig. 2H). We observed that clusters enriched in young mice showed high degree of correspondence with the established KP modules compared to clusters mostly prevalent in old mice (Fig. 2H). For example, the young age-specific cluster 1 matches the AT1-like state identified in the KP GEMM (Fig. 2H and I), which has been recently shown to be a driver of drug resistance in the context of the autochthonous model(*22*). Cluster 6 recapitulated several common states of KP evolution, including the high plasticity, mixed, and gastric-like states (Fig. 2H and I), while also expressing higher levels of epithelial lineage genes such as Sftpc, Epcam, Lyz2 and Nkx2-1 (fig. S4B and table S2). In contrast, clusters 7 and 8, which also arise through the canonical path of KP evolution, more predominantly match the mesenchymal states that evolve within the autochthonous model(*22*) (Fig. 2H and I). However, their proportions significantly differ depending on the age of the lung, with cluster 7 being more predominantly found in LUAD developing in the young lung, and cluster 8 in the old lung (Fig. 2E). Strikingly, old age also drove the evolution of terminal branches 4 and 5 from cluster 2 directly (Fig. 2D to G). Intriguingly, these cell populations did not resemble canonical KP clusters (Fig. 2H) nor did they strongly match any other cell type or lineage signatures in the body(*23–25*) (fig. S5), suggesting that LUAD developing in an aged lung deviates away from canonical KP evolution. Collectively, these data demonstrate that the age of the host environment shapes the evolution of KRAS-driven LUAD and support a model of tumor progression in which the age of the TME plays a key role in the selection of either a canonical or non-canonical evolutionary path, leading to the expansion of different stable cell populations (Fig. 2J).

To gain insights into the properties of LUAD cells that arise in the old lung environment, we performed GSEA of differentially expressed genes in malignant cells between young and old mice based on Canonical Pathways (CP) and Gene Ontology (GO) (Fig. 3A and B, tables S3 to 5). Strikingly, these analyses unveiled a common theme, stemness, as evidenced by pathways associated with pluripotency, cell fate commitment, and specification (Fig. 3A to C, table S5). Indeed, some of the most highly differentially expressed genes between LUAD cells developing in old compared to young mice were well-known factors involved in cancer stemness such as *Sox9*(*26*) (Fig. 3D and E), *Cd44*(*27*) (Fig. 3F and G), *Sox4*(*28*)*, Zeb2*(*29*) *and Hmga2*(*30*) (fig S4B, fig S6A and table S3). Strikingly, while these features are highly predominant in the cell populations that evolve through the non-canonical path (clusters 4 and 5), they are also a feature of cluster 8, which evolves across the canonical LUAD evolutionary continuum but is more predominant in old age (Fig. 3D, F and H, fig. S4B and S6A), highlighting the acquisition of stemness as a major feature of LUAD that evolves in the context of the old lung regardless of the specific path of evolution. Interestingly, clusters 4 and 5, but not cluster 8 which develops through the canonical evolutionary path, displayed an absence of defining markers apart from stemness-related genes (fig. S5, fig S6A and table S2) and a low number of detected genes and reads (fig. S6B), despite having high cellular complexity (fig. S6B). This observation was reminiscent of transcriptional quiescence, a staple of deeply quiescent stem cells(*31*), which was corroborated by the reduced numbers of LUAD cells with high RNA-Pol II CTD-Ser5, a marker of active transcription, in LUAD tumors developing in old mice (fig. S6C and D). This suggested that the aging-specific non-canonical path of LUAD evolution converges on a primitive stem-like state. In line with this idea, clusters 4 and 5 also displayed a much more pronounced loss of genes associated with lineage identity of alveolar type II (AT2) cells, the cells of origin of LUAD(*32*), including *Sftpc*, *Nkx2-1*, *Lyz2* (Fig. 3H and I, fig. S4B), indicating that old age accelerates the loss of lineage fidelity characteristic of LUAD cells(*15*). Together, this data supports a model whereby LUAD cells present in the old lung diverge from canonical LUAD evolution previously described in the context of young age(*15, 21*). Instead, when developing in an old environment, LUAD cells evolve towards a unique primitive stem-like state characterized by enhanced stemness and accelerated loss of lineage fidelity (Fig. 3J).

**Figure 3.**
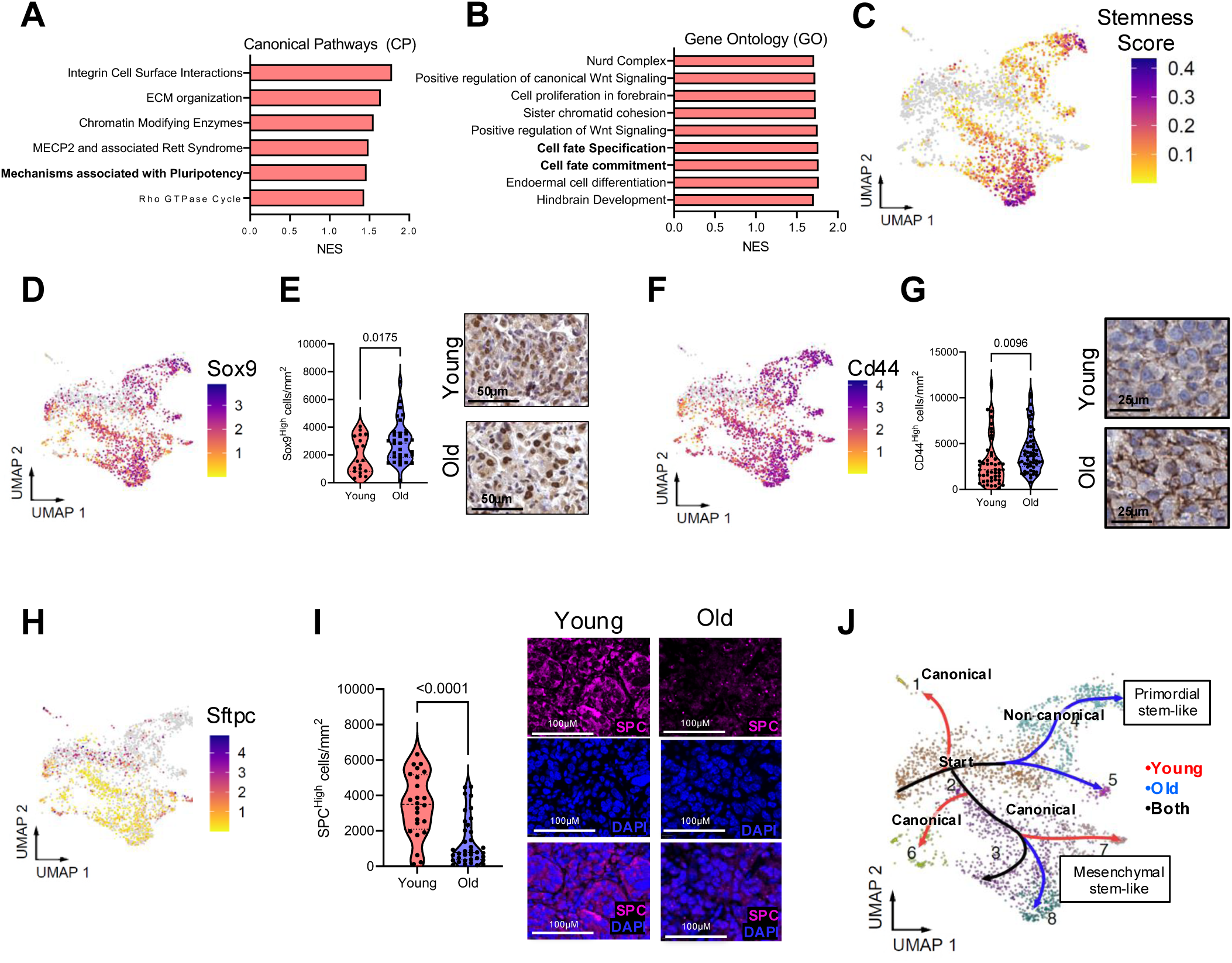
Aging drives LUAD’s differential evolution towards a primordial stem-like state. **A.** GSEA analysis of Canonical Pathways of epithelial malignant cells isolated from old compared to young mice. **B.** GSEA analysis of Gene Ontology terms of epithelial malignant cells isolated from old compared to young mice **C.** Stemness score generated from genes in bolded CP and GO pathways from A and B plotted onto the UMAP of epithelial malignant cells **D.** Log2(normalized gene expression) for Sox9 plotted onto the UMAP of epithelial malignant cells **E.** Quantification of the number of Sox9^High^ cells in young (5-6 month) and old (22-24 month) KP tumor-bearing mouse lungs along with representative IHC images (n=19 tumors from 4 young mice, n=33 tumors from 4 old mice, t-test) **F.** Log2(normalized gene expression) for CD44 plotted onto the UMAP of epithelial malignant cells **G.** Quantification of the number of CD44^High^ cells in young (5-6 month) and old (22-24 month) KP tumor-bearing mouse lungs along with representative IHC images (n=49 tumors from 5 young mice, n=57 tumors from 5 old mice, t-test) **H.** Log2(normalized gene expression) for Sftpc plotted onto the UMAP of epithelial malignant cells **I.** Quantification of the percentage of SPC^High^ cells in young (5-6 month) and old (22-24 month) KP tumor-bearing mouse lungs along with representative IF images (n=24 mice tumors from 5 young mice, n=37 tumors from 5 old mice, t-test). **J.** UMAP of epithelial malignant cells with consensus arrows depicting evolutional trajectory from Figure 2 and boxes to characterize the age-related canonical and non-canonical terminal branches. Dotted lines in violin plots represent the median and quartiles.

The TME is now recognized as a dynamic interaction arena, in which tumor cells interact with both resident and recruited host cells sharing and/or competing for soluble growth factors, cytokines, chemokines, nutrients, metabolites and extracellular matrix components(*33*). Accordingly, the identity, number and function of cellular components of the TME radically influences the fitness landscapes that are selected for in tumor cell subpopulations, as well as the niche-specific signaling that supports or inhibits the survival, growth, expansion and ability to metastasize of tumor subpopulations(*34*). Having found that the old lung environment significantly alters the fate of otherwise similar LUAD cells, next we asked which factors within the aged TME may be steering this differential LUAD evolution. In-depth analysis of our scRNA-Seq data focusing on TME cells (fig. S7) identified 13 other major cell types which can be further classified into 40 different cell subtypes present in LUAD’s TME (fig. S7 to S12). While the aged TME showed moderate alterations in the overall proportions of major cell populations compared to young counterparts, such as in NK cells, CD8^+^ T cells, neutrophils, and fibroblasts (fig. S7E), the biggest shift caused by old age was observed in the distribution of different cell subsets within these major populations (fig. S8E and S9F). For instance, we detected an enrichment of CTHRC1^+^ fibroblasts within the aged TME (fig. S8D to H), which are defined by tenascin C expression (fig. S8B) and are known to be pro-fibrotic and associated with human pulmonary diseases(*35*), possibly contributing to altered tissue architecture and tumor progression. Within the immune compartment, we noted an age-induced accumulation of NK-like CD8^+^ T-cells (fig. S9C to I), which have been previously reported to accumulate in aged individuals(*36*) and during cancer progression(*37*), and have been proposed as a potential therapeutic target(*38*). This suggests that aging primarily influences the TME’s cell phenotypic diversity (tables S6 and S7) rather than causing dramatic changes in the abundance of major cell types.

Considering the age-imposed phenotypic changes observed in many major cell types of the TME, we reasoned that these differences likely translate into distinct niche signaling mechanisms of communication between the TME and tumor cells thereby steering cell fate decisions and consequently altering LUAD’s features. Quantitative inference and analysis of intercellular communication networks from our scRNA-Seq data using CellChat(*39*) unexpectedly revealed that normal, non-malignant, epithelial cells, while low in abundance (fig. S7E), dominated the niche signaling within the aged LUAD TME, with the most numerous and strongest interactions (Fig. 4A). Deconvolution of the epithelial normal compartment of the TME, revealed 3 distinct subpopulations, an AT2 population, a Club/Ciliated population, and an epithelial proliferative population that did not match any of the canonical epithelial cell types of the lung (Fig. 4B, fig. S12A and B, table S8). Interestingly, it is this epithelial proliferative population that changes the most with old age, not only increasing significantly in proportion within the aged TME (Fig. 4C), but also becoming the most interactive population with a pronounced change in both outgoing and incoming interaction strengths when compared with the young TME (Fig. 4D, fig. S12C to F, table S9). In the alveoli, AT2 cells maintain lung homeostasis and enable re-generation after injury by proliferating and differentiating into new AT1 cells specialized for gas exchange(*40*). Recently an AT2-lineage population of cells, ADIs, were shown to arise during alveolar regeneration as an intermediate state between activated AT2 cells and their differentiation into AT1 cells, characterized by the absence of both AT2 and AT1 markers(*41, 42*). ADIs are rare at steady state but are significantly induced after injury by IL-1-mediated inflammatory signaling(*41*). ADIs aberrantly accumulate in conditions of continued damage or chronic inflammation in both mice and humans, thereby preventing differentiation into AT1 cells and impairing alveolar regeneration(*41, 42*). Considering our scRNA-Seq data as well as the well-known effects of aging in promoting damage (e.g., fibrosis) and chronic inflammation in the lung(*43*), we reasoned that the epithelial proliferative population, which arises in the context of old age and makes up more than 50% of the epithelial normal compartment in the old lung TME (Fig. 4C), might be ADIs. ADIs are characterized by the expression of senescence marker genes and p53 as well as induction of G2/M signaling while maintaining a proliferative capacity marked by the expression of Ki67 and upregulated glycolysis(*41, 42*). GSEA of the differentially expressed genes in the normal epithelial proliferative subpopulation when compared to the remaining epithelial normal cell populations revealed similarities between this population and ADIs, as observed by the enrichment of pro-proliferative gene signatures (e.g., myc and E2F targets, mTORC1 signaling) as well as gene signatures related to p53 and senescence induction (e.g., G2M checkpoint, DNA repair; Fig. 4E and table S10). In line with this, the epithelial proliferative population identified in our scRNA-Seq largely matched the described ADI transcriptional signature(*41, 42*) (Fig. 4F). IF analysis validated the presence of normal AT2 (SPC^+^/ p53^+^) cells within the tumors and showed a decrease in AT2 cells within the tumors of old mice (Fig. 4G and fig. S13A) corroborating the scRNA-Seq analysis (Fig. 4C). This age-induced change in AT2 cells was accompanied by an increased number of AT1 cells within LUAD tumors (Fig. 4H and fig. S13B). The presence of ADIs (KRT8^+^/p53^+^ cells (*41, 42*)) and their increase in LUAD tumors from old mice was also validated by IF (Fig. 4I). Importantly, while we also observed a substantial increase in ADIs in the normal lung tissue immediately adjacent to LUAD in the context of old age (Fig. 4J), in non-tumor bearing (NTB) lungs the abundance of ADIs was much reduced and comparable between young and old mice (fig. S13C). This suggests that the presence of LUAD interacts with the old alveolar epithelium to drive ADI accumulation within the tumor and the adjacent lung tissue. To further validate this phenomenon and better distinguish between host-derived and transplanted LUAD cells, we employed a genetic lineage tracing approach. We transplanted KP cells into Spc-Cre/Kb^Strep^ mice, which express a Cre-driven StrepTagII(*44*) that is permanently incorporated into the MHC-I molecules of SPC-expressing AT2 cells (Fig. 4K). By immunostaining for both Krt8 and StrepTagII in lungs from these mice, we confirmed the presence of Krt8-expressing cells that were indeed derived from host SPC-expressing AT2 cells (Krt8^+^/StrepTagII^+^) (Fig. 4K). In line with our previous observation, this analysis showed an age-related increase in host-derived ADIs (Fig. 4L and M) further confirming ADI accumulation as a feature of the old LUAD TME (Fig. 4I to M), but not of NTB old lungs (fig. S13C).

**Figure 4.**
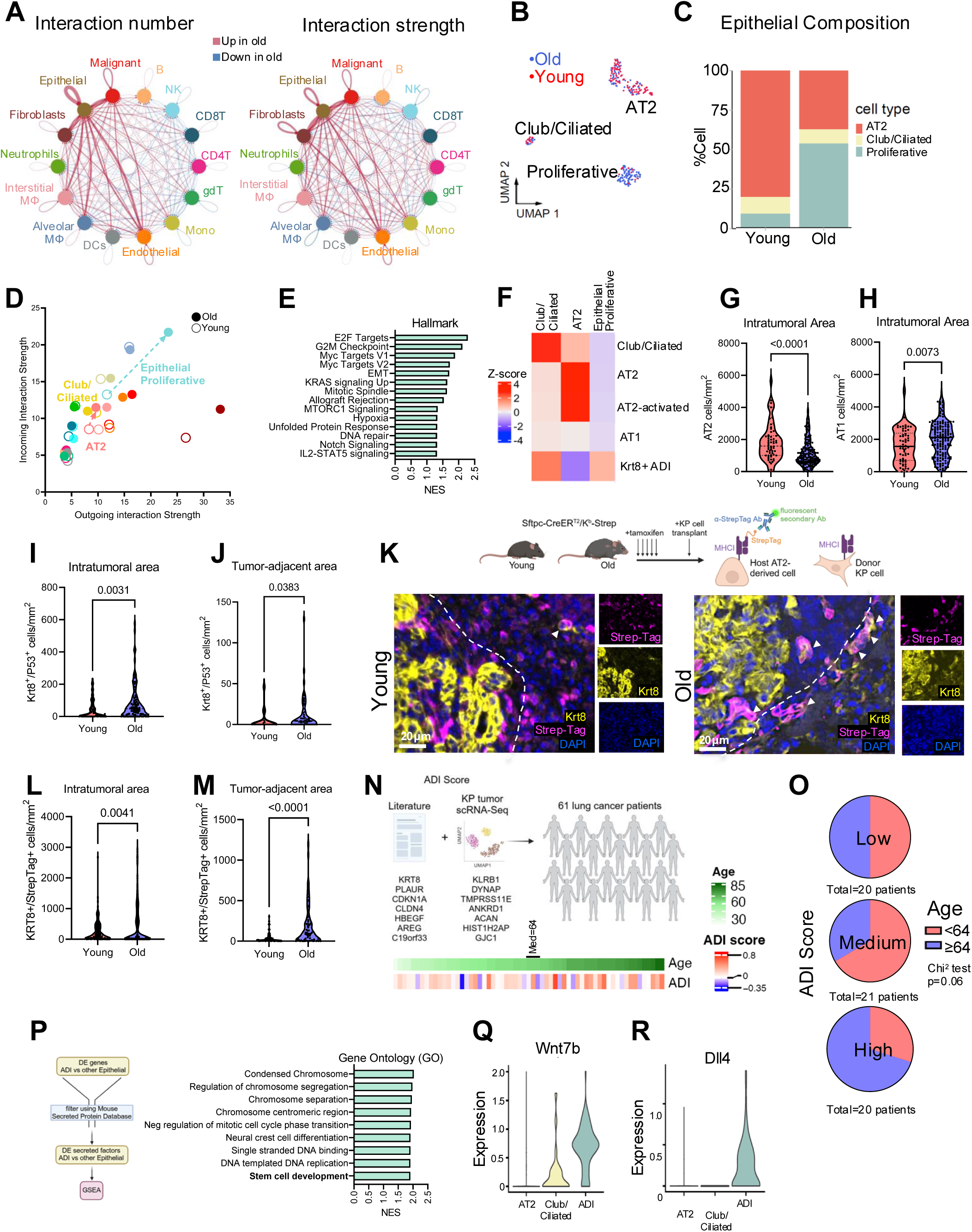
Age-related TME communication is mediated largely by Alveolar Differentiation Intermediate Cells. **A.** Chord diagrams from interaction number and strength based on CellChat analysis from cells within the TME cells from KP tumors isolated from young and old mice. Red and blue lines represent interactions that are higher and lower in old tumors, respectively, compared to young tumors. **B.** UMAP projection of normal epithelial cells, colored by age. **C.** Bar graph of normal epithelial cell type proportions in KP tumors from young and old mice **D.** Scatterplot showing the incoming and outgoing interactions strength of cell types within the young (open circles) and aged (solid circles) KP TME **E.** Hallmark GSEA analysis of top 15 most enriched pathways in epithelial proliferative cells compared to other normal epithelial cells. **F.** Heat map showing z-scores for similarity of normal epithelial subclusters with signatures of known epithelial cell types. **G.** Quantification of AT2 cells in tumors from young and old mice, defined as Spc^+^/P53^+^ cells or Spc^+^/Pdpn^-^/Krt8^-^ cells (n=62 tumors from 5 young mice, n=162 tumors from 5 old mice, t-test) **H.** Quantification of AT1 cells in tumors from young and old mice, defined as Pdpn^+^/P53^+^ cells or Pdpn^+^/Spc^-^/Krt8^-^ cells (n=62 tumors from 5 young mice, n=162 tumors from 5 old mice, t-test) **I.** Quantification of Krt8^+^/P53^+^ ADI cells in tumors from young and old KP tumor-bearing mice (n=43 tumors from 6 young mice, n=97 tumors from 5 old mice, t-test) **J.** Quantification of Krt8^+^/P53^+^ ADI cells in tumor-adjacent areas (200µm radius around tumor perimeter) from young and old KP tumor-bearing mice (n=26 tumor-adjacent areas from 6 young mice, n=47 tumor-adjacent areas from 5 old mice, t-test) **K.** Schematic of Spc-Cre/Kb-Strep mouse transplantation experiment (top), and representative images of IF staining for Krt8, StrepTagII and DAPI in young (5-6 month) and old (22-24 month) KP tumor-bearing mouse lungs (bottom) **L.** Quantification of Krt8^+^/StrepTagII^+^ ADI cells in tumors from young and old KP tumor-bearing mice (n=5 mice per group, n=207 tumors from old mice, n=179 tumors from young mice, t-test). Dotted lines represent tumor borders. **M.** Quantification of Krt8^+^/StrepTagII^+^ ADI cells in tumor-adjacent areas (200µm radius around tumor perimeter) from young and old KP tumor-bearing mice (n=5 mice per group, n=80 tumor-adjacent areas from old mice, n=69 tumor-adjacent areas from young mice, t-test) **N.** Heatmap showing z-score for the ADI signature (genes depicted in upper schematic) in 61 lung cancer patients, ordered by age. Black line represents the median age. **O.** Pie charts showing the distribution of 61 lung cancer patient samples (separated by median age=64) within different bins of ADI scores (low<0.04, medium=0.04-0.14, high ≥0.14, Chi^2^ test, p= 0.06334). **P.** GSEA analysis using Gene Ontology (GO) terms of differentially expressed genes in ADIs compared to other epithelial normal cells, after filtering 2 for secreted factors using a Mouse Secreted Protein Dataset(*85*) **Q.** Violin plot showing the log2(Normalized gene expression) for Wnt7b in epithelial subtypes from KP tumor-bearing mice. **R.** Violin plot showing the log2(Normalized gene expression) for Dll4 in epithelial subtypes from KP tumor-bearing mice. Dotted lines in violin plots represent the median and quartiles.

Recently ADI-like states have been implicated in the transformation of AT2 cells into lung cancer(*25, 45*). However, the role of LUAD-infiltrating ADIs remains unknown. Thus, we questioned whether ADIs are also present and may also be relevant in human cancers. Therefore, we generated an ADI signature score composed of established marker genes from the literature(*46*) as well as non-overlapping marker genes from our own dataset within the top 10 markers of the epithelial proliferative population (Fig. 4N and table S11). We then leveraged a scRNA-Seq database of human LUAD samples(*47*), and uncovered the presence of cells in human samples matching to our ADI score (Fig. 4N), with a trend towards an increase in these cells with age (Fig. 4O). This finding is concordant with a previous report demonstrating the existence of these cells in humans with idiopathic pulmonary fibrosis (IPF)(*46*) and further establish the relevance of our findings for human disease. Interestingly, ADIs interact with most of the different LUAD clusters (fig. S12C and S13D), supporting that ADIs ability to steer LUAD evolution is likely driven by their abundance rather than a specific interaction with a subset of LUAD cells. Consistently, our analyses also revealed that a large part of the niche signaling originating from by ADIs is mediated by soluble or secreted ligands (fig. S13E, tables S9 and S12), suggesting that ADIs may not only signal to the LUAD cells in their vicinity but also have the potential to affect distal LUAD cells and highlighting the importance of ADIs not only within the TME, but also in the adjacent normal lung. Supporting old age-induced ADI accumulation in directing the differential evolution of LUAD into a primordial stem-like state, analyses of the cell-to-cell communication mechanisms between ADIs and LUAD cells revealed several of these factors are well-known regulators of stemness, such as Wnt and Notch signaling (Fig. 4P to R, fig. S13D and F, tables S9 and S12) with ADI-derived ligands such as Wnt5a, Wnt7b, Dll4 and Jag1 found to interact accordingly with Fzd and Notch receptors on malignant cells (Fig. 4P to R, fig. SD and F, tables S9 and S12).

Taken together, these findings demonstrate for the first time a role for the aged normal lung alveolar epithelium in shaping LUAD evolution toward a primordial stem-like state, which is typically associated with poor prognosis(*48–50*), putting the aging process center stage in dictating LUAD’s features. We traced this phenomenon, at least in part, to the ability of old age to prime the alveolar epithelium for ADI accumulation upon the presence of LUAD, which in turn creates a niche rich in stemness-inducing signals. Recent publications have highlighted hematopoietic aging as a major regulator of LUAD progression through an IL-1-mediated mechanism(*51*), as well as the reduced stemness of AT2 cells with old age, which lose their ability to differentiate into AT1 cells as well as their ability to initiate LUAD(*18*). Considering that ADIs are induced by the IL-1/IL1R1 signaling axis(*52*), it is plausible that the increase in IL-1 produced by the aged immune system recruited into the LUAD TME(*51*) drives AT2 cell differentiation, leading to the accumulation of cells stuck in the intermediate ADI state due to their age-induced inability to transition into AT1 cells(*18*). Additional work will be necessary to tease apart the mechanisms by which old age causes ADI accumulation. How broadly applicable might these finds be? Our findings that the normal lung alveolar epithelium is an important component of LUAD’s TME was only possible due to the choice of the model used, a transplant-based model that does not genetically engineer AT2 cells within the lung, thus it is likely that our findings will lead to the discovery of additional factors that cause ADI accumulation but were previously not detected (e.g. smoking, lung infections). Moreover, the lung is a major site of metastasis for several different cancers, including cancers of major incidence like breast cancer and melanoma. Thus, it is possible that ADI accumulation in the old lung environment also occurs during metastatic homing to this tissue which has potential important consequences for prognosis. Moreover, this comprehensive analysis of LUAD and its TME by age at single cell resolution provides a valuable resource for understanding age-related differences and provide a blueprint for the discovery of new therapeutic targets for LUAD in the context of old age, thereby having the potential to inform the development of age-specific treatment strategies.

## Methods

### Mice

Wild-type C57BL/6J mice of different ages were obtained from the National Institutes of Aging, NIH. Mice were housed in an AAALA-accredited animal facility in ventilated pathogen-free cages with food, water, bedding and nestling materials. All experiments involving mice were approved by the Institutional Animal Care and Use Committee (IACUC) prior to initiation. To induce Cre recombination and Strep-TagII expression in Spc-expressing cells, Spc-Cre/Kb^Strep^ mice were administered 75 mg tamoxifen (Sigma, 10540-29-1) per kg of body weight via intraperitoneal (IP) injection, once daily over 5 consecutive days. Histology or KP transplantation experiments were performed 10-14 days after the last tamoxifen injection.

### Cell culture

KP 1.9 cells(*53, 54*) (gift from Dr. Zippelius) were maintained in DMEM high glucose medium (Cytiva) supplemented with 10% FBS (Gibco) and 1% penicillin/streptomycin (Cytiva) at 37°C and 5% CO_2_.

### Syngeneic Transplantation Model of LUAD

KP 1.9 cells were counted using a Countess II FL automatic hemocytometer (Invitrogen) and collected in PBS at a concentration of 1 x 10^6^ cells/ml. Mice were restrained using a conical tube and their tails were warmed with hot water to improve tail vein visibility. 100 µl of cell suspension (1 x 10^5^ cells) was injected into the tail vein using a 27G needle (BD Biosciences). Four weeks post-injection, mice were euthanized via CO_2_ inhalation followed by cervical dislocation, and the lungs were collected for subsequent experiments.

### Immunohistochemistry

Mouse lung tissue samples were dissected and fixed in 10% formalin. After 24 hours, samples were transferred to 70% ethanol for storage. Paraffin-embedded tissue cassettes were prepared, cut into sections, and stained using hematoxylin & eosin or Masson’s Trichrome stains by IDEXX BioAnalytics (Westbrook, ME, USA). Unstained slides were deparaffinized and rehydrated by sequential incubations in a series of alcohol solutions comprised of 2x xylene for 10 minutes, 2x 90% ethanol for 3 minutes, 2x 70% ethanol for 3 minutes, 2x 50% ethanol for 3 minutes, and 2x diH_2_0 for 1 minute. Antigen retrieval was performed by heating under pressure for 15 minutes in citrate buffer (10 mM citric acid, 0.05% Tween 20, pH 6.0). Samples were transferred to 3% H_2_O_2_ in water for 5 minutes, rinsed in TBS-T, and then permeabilized for 15 minutes in 0.5% Triton X-100 in PBS. Samples were then rinsed in TBS-T, followed by blocking for 1 hour in 3% BSA (m:v)/ 10% goat serum (v:v) and/or 10% horse serum (depending on the host species of antibodies to be used in each panel) in TBS. Slides were then incubated with the primary antibodies: rabbit anti-Sox9 (Millipore Cat# AB5535, RRID:AB_2239761); rat anti-CD44 (Thermo Fisher Scientific Cat# 14-0441-82, RRID:AB_467246); rabbit anti-SPC (Seven Hills Bioreagents Cat# WRAB-76694, RRID:AB_2938817); rabbit anti-RNA polymerase II CTD phospho S5 (Abcam Cat# ab5131, RRID:AB_449369) diluted in 3% BSA (m:v)/ 10% goat serum (v:v) and/or 10% horse serum (depending on the host species of antibodies to be used) in TBS in a humid container at 4°C overnight. The next day, slides were washed 2x 5 minutes in TBS-T and incubated for 30 minutes at RT with ImmPRESS HRP anti-rabbit (MP-7451-15, Vector Laboratories) or anti-rat polymer detection reagent (MP-7444-15, Vector Laboratories). Samples were then washed 2x 5 minutes in TBS-T and developed using the ImmPACT DAB HRP Substrate Kit (Vector Laboratories), followed by counterstaining with hematoxylin (Vector Laboratories). Slides were then washed 2x for 1 minute each in 100% isopropanol, mounted using Vectamount Express Mounting Medium (Vector Laboratories, H-5700-60), and dried at room temperature overnight before imaging on an Aperio imager (Leica Biosystems, Nussloch, Germany) at 20x magnification.

### Tissue immunofluorescence

Mouse lung tissue samples were dissected and fixed in 10% formalin. After 24 hours, samples were transferred to 70% ethanol for storage. Paraffin-embedded tissue cassettes were prepared and cut into sections by IDEXX BioAnalytics (Westbrook, ME, USA). Slides were deparaffinized and rehydrated by running them through an alcohol series of 2x xylene for 10 minutes, 2x 90% ethanol for 3 minutes, 2x 70% ethanol for 3 minutes, 2x 50% ethanol for 3 minutes, and 2x diH_2_0 for 1 minute. Antigen retrieval was performed by heating under pressure for 15 minutes in Citrate buffer (10mM citric acid, 0.05% Tween 20, pH 6.0) or Tris-EDTA buffer (10 mM Tris, 1 mM EDTA, 0.05% Tween 20, pH 8.0), depending on primary antibody recommendations. Samples were rinsed in TBST and then permeabilized for 15 minutes in 0.5%Triton X-100 in PBS. Samples were then rinsed in TBST followed by blocking for 1 hour in 3% BSA (m:v)/ 10% goat serum (v:v) and/or 10% horse serum (depending on the host species of antibodies to be used) in TBS. Slides were then incubated with the primary antibodies : rabbit anti-SPC (Seven Hills Bioreagents Cat# WRAB-76694, RRID:AB_2938817) ; rat anti-KRT8 (DSHB Cat# TROMA-I, RRID:AB_531826); goat anti-Pdpn (R and D Systems Cat# AF3244, RRID:AB_2268062); mouse anti-P53 (Proteintech Cat# 60283-2-Ig, RRID:AB_2881401); rabbit anti-RNA polymerase II CTD phospho S5 (Abcam Cat# ab5131, RRID:AB_449369); goat anti-Pdgfrα (R and D Systems Cat# AF1062, RRID:AB_2236897); mouse anti-Tenascin C (Thermo Fisher Scientific Cat# MA5-16086, RRID:AB_11152811); rabbit anti-CD8 AF488 (Abcam Cat# ab237364, RRID:AB_2940904); rabbit anti-NKG2A (Bioss Cat# bs-2411R, RRID:AB_10860210), rabbit anti-Strep-tag II(Abcam Cat# ab307676, RRID:AB_3674753) diluted in 3% BSA (m:v)/ 10% goat serum (v:v) and/or 10% horse serum (depending on the host species of antibodies to be used) in TBS in a humid container at 4°C overnight. The next day, slides were washed 2x 5 minutes in TBS-T and incubated for 1 hour at RT with secondary fluorescence antibodies: rabbit anti-goat IgG AlexaFluor 488 (Thermo Fisher Scientific Cat# A-11078, RRID:AB_2534122); donkey anti-rat IgG AlexaFluor 555 (Thermo Fisher Scientific Cat# A78945, RRID:AB_2910652); donkey anti-mouse IgG IRDye 680RD (LI-COR Biosciences Cat# 926-68072, RRID:AB_10953628); donkey anti-Rabbit IgG CF750 (Sigma-Aldrich Cat# SAB4600372-50UL, RRID:AB_3674754); goat anti-Rabbit IgG AlexaFluor 594 (Thermo Fisher Scientific Cat# A32740, RRID:AB_2762824). The secondary antibody solution was then replaced with DAPI (0.2 µg/ml) in PBS and incubated for 10 minutes before mounting using VECTASHIELD Antifade Mounting Medium with DAPI (Vector Laboratories), and coverslips were sealed with nail polish. Slides were imaged using an Akoya Biosciences HT system (Akoya Biosciences) at 20x.

### QuPath analysis

Whole slide ScanScope Virtual Slide (SVS) image files were imported into QuPath(*55*) (version 0.4.1), and tumors were annotated using the wand tool. For IHC analyses, cell detection was performed using either optical density sum or hematoxylin (depending the presence or absence of hematoxylin counterstain) to detect all cells with a 10 µm nuclear size minimum threshold and a 5 µm cell expansion radius when profiling a non-nuclear (cytoplasmic or membrane) stain. Positive cell detection was used using a single positivity threshold using the following thresholds for the corresponding stains: P16, 0.4 (DAB); p-H2A.X, 0.35 (DAB); Sox9, 0.5 (DAB); CD44, 0.25 (DAB); SPC, 37 (fluorophore); RNA Pol II CTD-pSer5, 0.45 (DAB); CD8, 27 (fluorophore); NKG2A, 32 (fluorophore); Pdgfrα, 8.5 (fluorophore); Tnc, 28 (fluorophore); Pdpn, 45 (fluorophore); Krt8, 19 (fluorophore); P53, 22 (fluorophore). For immunofluorescence cell typing analyses, a single measurement classifier was performed for each color, with the positivity threshold selected based on the signal intensity histogram and using a secondary-only negative control as a guide. All single object classifiers were then compiled into one composite classifier to obtain the number of cells under each combination of fluorescence markers.

### Tumor histological grading

Whole slide image files of hematoxylin & eosin-stained mouse lungs were analyzed using GLASS-AI(*14*) v1.1 (https://github.com/jlockhar/GLASS-AI). GLASS-AI is a purpose-built tool for automatically analyzing mouse models of LUAD and uses the tumor grading conventions developed in the Tyler Jacks laboratory(*55*). Regions of stromal desmoplasia, which correspond to Grade 5 and are not recognized by GLASS-AI, were manually annotated and integrated with the GLASS-AI output using the GLASS-AI Annotation Editor (https://github.com/jlockhar/GLASS-AI-annotation-editor). Individual tumors were assigned overall grades based on the highest morphological grade present that comprised at least 10% of the tumor’s area. The results of these analyses were summarized in R using GLASS-AI ReportR (https://github.com/jlockhar/GLASS-AI-ReportR).

### Flow Cytometry

Mice were euthanized via CO_2_ inhalation followed by cervical dislocation, and lungs were collected in PBS on ice. Lungs were minced using scissors and a razor blade until no visible chunks remained. Minced lungs were transferred to a 5 ml epitube containing 4ml of dissociation buffer composed of 2 mg/ml Collagenase I (Worthington), 0.5 mM CaCl_2_ in DMEM/F12 1:1 media (Corning). Samples were placed on a rotor in a 37°C incubator for 30 minutes. Undigested tissue was spun down at 300xg for 45 seconds. The supernatant of released cells was transferred to a tube containing 5 ml of FBS on ice to inactivate the dissociation enzymes. The undissociated tissue was triturated using a P1000 pipette and 2.5 ml of fresh dissociation buffer was added to the remaining tissue. The samples were dissociated once more for 30 minutes. Cells from both dissociations were combined and passed through a 70 µm cell strainer. The blunt end of the inner plunger of a 5 ml syringe was used to force any remaining tissue pieces through the strainer. The filtered sample was centrifuged at 300xg for 7 minutes. The cell pellet was resuspended in PBS and stained for viability with LIVE/DEAD Fixable Near-IR Dead Cell Stain (Life Technologies) for 15 minutes at room temperature, washed in PBS, and spun down at 300xg for 7 minutes. Cells were then resuspended in 100µl PBS with Trustain FcX PLUS (anti-mouse CD16/32) (BioLegend), True-Stain Monocyte Blocker (BioLegend), and Super Bright staining buffer (eBioscience) for 10 minutes before adding in the following antibody cocktails and incubating for 15 minutes at room temperature: <u>immune cell panel</u>: rat anti-CD3-BUV395 (BD Biosciences Cat# 740268, RRID:AB_2687927), rat anti-CD11b-BV785 (BioLegend Cat# 101243, RRID:AB_2561373), hamster anti-CD11c-BV605 (BioLegend Cat# 117333, RRID:AB_11204262), rat anti-IA/IE-BV650 (BioLegend Cat# 107641, RRID:AB_2565975), rat anti-CD24-BV421 (BioLegend Cat# 101825, RRID:AB_10901159), rat anti-Ly6G-BV510 (BioLegend Cat# 127633, RRID:AB_2562937), rat anti-CD25-BB515 (BD Biosciences Cat# 564424, RRID:AB_2738803), rat anti-Ly6C-PerCP-Cy5.5 (BioLegend Cat# 128028, RRID:AB_10897805), rat anti-CD45-PE (BioLegend Cat# 103105, RRID:AB_312970), rat anti-Gr1-Pe/Dazzle594 (BioLegend Cat# 108452, RRID:AB_2564249), mouse anti-CD64-PECy7 (BioLegend Cat# 139313, RRID:AB_2563903), rat anti-CD8a-AF647 (BioLegend Cat# 100724, RRID:AB_389326), rat anti-CD4-R718 (BD Biosciences Cat# 566939, RRID:AB_2869957); <u>stromal cell panel</u>: rat anti-EpCAM-BV421 (BioLegend Cat# 118225, RRID:AB_2563983), rat anti-CD90-BV605 (BioLegend Cat# 105343, RRID:AB_2632889), rat anti-IA/IE-BV650 (BioLegend Cat# 107641, RRID:AB_2565975), rat anti-CD146-FITC (BioLegend Cat# 134706, RRID:AB_2143525), rat anti-CD45-PE (BioLegend Cat# 103105, RRID:AB_312970), rat anti-CD31-PECy7 (BioLegend Cat# 102417, RRID:AB_830756), rat anti-CD34-R718 (BD Biosciences Cat# 567304, RRID:AB_2916545). Samples were washed with PBS containing 2%FBS and then resuspended in PBS for analysis using a BD LSR II Flow cytometer (BD Biosciences) running BD FACSDiva software (BD Biosciences). After data collection, flow data was analyzed using FlowJo software (version 10.8.1, BD Biosceinces). Gating strategies were adapted from Hasegawa K. et al.(*56*) and Yu YRA, et al.(*57*).

### Bronchioalveolar Lavage

Mice were anesthetized using xylene/ketamine solution following institutional guidelines. An incision was created to expose the trachea, and a catheter was inserted into the trachea. A syringe loaded with 1 ml of PBS + protease inhibitor solution (5 µg/ml Leupeptin, 1 µg/ml Aprotinin, 0.5 µg/ml Pepstatin A; Research Products International) was injected via the catheter into the trachea and aspirated back to collect bronchioalveolar lavage fluid (BALF). This lavage was repeated for a total of 5 times to recover approximately 5 ml of total BALF. Mice were then euthanized via cervical dislocation. BALF samples were centrifuged at 300xg for 5 minutes to remove any cells. The BALF supernatant was then concentrated into powder form using a Speed vac at room temperature and resuspended in 500 µl of PBS + protease inhibitor solution. One-third (166 µl) of this solution was utilized to run the Proteome Profiler Mouse XL Cytokine Array (R&D Systems) following the manufacturer’s instructions, followed by imaging using an Odyssey LI-COR Imager.

### Bulk RNA-Seq

RNA was extracted from cultured KP cells (from 3 different passages) using the RNeasy Mini Kit (Qiagen). RNA samples were sent to Novogene Corporation Inc. (Sacramento, CA) for RNA-Seq library preparation and sequencing. A total of 140,783,878 reads were generated by a NextSeq 500 with a 151-base paired-end read run. Adapter sequences were trimmed using Novogene’s in-house Perl scripts, and the raw reads were then aligned to the mouse genome (mm39) using Hisat2(*58*) v2.0.5. Gene expression was assessed as read count at the gene level with featureCounts(*59*) v1.5.0-p3. Fragments Per Kilobase of transcript per Million mapped reads (FPKM) for each gene were calculated for the correlation analysis between scRNA-seq epithelial malignant clusters and bulk RNA-seq samples. The FPKM values were later used to calculate correlation with scRNAseq data.

### Single-Cell RNA-Sequencing sample preparation

KP tumors were micro-dissected using scissors and fine tip forceps from lungs collected from tumor-bearing mice and transferred to a 6cm petri dish. Tumors were minced into a fine slurry using scissors and a razor blade, and transferred to a 5ml tube containing 2.5 ml of digestion buffer from the Miltenyi mouse tumor dissociation kit (Miltenyi Biotec, Cat# 130-096-730). Tumors were placed on a rotor in a 37°C incubator. After 20 minutes of dissociation, samples were centrifuged at 300xg for 45 seconds to pellet undigested tissue, and the supernatant containing released cells was transferred to a tube containing 3 ml FBS on ice. 2.5 ml of digestion buffer was added back to the original tube and the remaining tissue was triturated using a P1000 pipette. After another 20 minutes of dissociation, both cell suspensions were combined and filtered through a 70 µm cell strainer. Cells were centrifuged for 7 minutes at 300xg, the supernatant was removed and cells were resuspended in 200-500 µl of PBS containing 0.04%BSA depending on the size of the cell pellet. Cells were counted using a manual hemocytometer and the concentration was adjusted to 1000 cells/µl in PBS containing 0.04% BSA for library preparation by the Molecular Genomics core at the Moffitt Cancer Center.

### Single-cell sequencing

Single-cell RNA sequencing was performed using the 10X Genomics Chromium System (10X Genomics, Pleasanton, CA) by the Molecular Genomics Core at the Moffitt Cancer Center. The cell viability and counts were obtained by AO/PI dual fluorescent staining and visualization on the Nexcelom Cellometer K2 (Nexcelom Bioscience LLC, Lawrence, MA). Cells were then loaded onto the 10X Genomics Chromium Single Cell Controller to encapsulate 7000 cells per sample. Single cells, reagents, and 10X Genomics gel beads were encapsulated into individual nanoliter-sized Gelbeads in Emulsion (GEMs), and reverse transcription of poly-adenylated mRNA was performed inside each droplet at 53°C. The cDNA libraries were then completed in a single bulk reaction by following the 10X Genomics Chromium NextGEM Single Cell 3’ Reagent Kit v3.1 user guide, and 50,000 sequencing reads per cell were generated on the Illumina NovaSeq6000 instrument. The Demultiplexing, barcode processing, alignment, and gene counting were performed using the 10X Genomics CellRanger v7.1.0 software. The results of the analysis were visualized using the 10X Genomics Loupe browser v6.4.1.

### Single-cell RNA-seq data processing, filtering, batch effect correction, and clustering

Raw sequencing reads from scRNA-seq were processed using Cell Ranger (v7.1.0, 10X Genomics). Briefly, the base call (BCL) files generated by Illumina sequencers were demultiplexed into fastq files based on the sequences of the sample index, and aligned against the GRCm38 mouse transcriptome using STAR(*60*). Cell barcodes and UMIs associated with the aligned reads were subjected to correction and filtering. Filtered gene-barcode matrices containing only barcodes with UMI counts passing the threshold for cell detection were imported to Seurat v4.0(*61*) for downstream analysis. Barcodes with fewer than 200 genes expressed or more than 10% UMIs originating from mitochondrial genes were filtered out; genes expressed in fewer than three barcodes were also excluded. This process resulted in 11,320 cells from three young KP mice, and 11,893 cells from three old KP mice. For each sample, standard library size and log-normalization were performed on raw UMI counts using *NormalizeData*, and the top 5000 most variable genes were identified by the “vst” method in *FindVariableFeatures*.

In each study, individual data were further integrated to remove batch effects using an anchor-based method(*62*) implemented in Seurat v4.0 using *FindIntegrationAnchors* and *IntegrateData* functions in Seurat with 8,000 “anchors” and top 40 principal components. Briefly, dimension reduction was performed on each data set using diagonalized canonical correlation analysis (CCA), and L2-normalization was applied to the canonical correlation vectors to project the datasets into a shared space. The algorithms then searched for mutual nearest neighbors (MNS) across cells from different datasets to serve as “anchors” which encoded the cellular relationship between datasets. Finally, correction vectors were calculated from “anchors” and used to integrate datasets.

From the integrated data, scaled z-scores for each gene were calculated using the *ScaleData* function in Seurat by regressing against the percentage of UMIs originating from mitochondrial genes, S and G2/M phase scores, and total reads count. A shared nearest neighbor (SNN) graph was constructed based on the first 40 principal components computed from the scaled integrated data. Louvain clustering (Blondel et al., 2008) was performed using the *FindClusters* function at resolution 1.2 for major cell type scRNA-seq data (15 clusters). Uniform manifold approximation and projection (UMAP) was used to visualize single-cell gene expression profile and clustering, using the *RunUMAP* function in Seurat with default settings. Differential expression analysis was performed using the *FindAllMarkers* function in Seurat with logfc.threshold=0.25, min.pct=0, and test.use=”wilcox”. Cells within each cluster were compared against all other cells. Genes with Bonferroni-corrected p-value < 0.05 and an average log-fold change > 0.25 and were considered differentially expressed. Clusters were annotated by comparing differential genes with canonical markers for major populations: B cells (*Cd79a, Cd79b, Cd19*), T cells (*Cd3e, Cd3d, Cd8a, Cd4*), NK cells (*Klrb1c, Ncr1, Nkg7*), Macrophages (*Cd68, Mrc1, C1qc, C1qb*), Monocytes (*Lyz2, Vcan, Chil3, Fn1*), Neutrophils (*S100a8, S100a9, Cxcl2*), cDC1 (*Xcl1, Cd36, Itgae*), cDC2 (*Itgax, H2-DMb1, Mgl2*), mregDC (*Fscn1, Ccl22, Cacnb3*), pDC (*Siglech, Ccr9, Bsl2*), Epithelial cells (*Sftpc, Krt7, Krt18*), Endothelial cells (*Pecam1, Cdh5*), Fibroblasts (*Col1a1, Col1a2*).

### Annotation of epithelial cells subtypes from scRNA-Seq data

Epithelial cells were extracted for further clustering analysis with resolution = 1.0 and yielded 11 clusters. Clusters 9-11 were annotated as normal epithelial cells based on their positive expression of KP mouse model genes (*Tp53*, *Kras*, and their targets(*63, 64*)), as well as the absence of tumor markers (*Krt7, Krt18, Krt19*). These three normal epithelial cells were further annotated as club/ciliated cells (*Tppp3, Tmem212, Dynlrb2)*, alveolar type 2 (AT2) cells (*Sftpc, Lyz2, Slc34a2*), proliferative cells (*Mki67, Cdk1, Ccna2*) respectively, based on previously reported markers(*65*) (fig. S3B). We also observed that proliferative cluster 11 co-express markers of Krt8+ alveolar differentiation intermediate (ADI) cells(*46*), such as *Krt8, Plaur, Cldn4, Areg*, and *Hbegf*. Clusters 1-8 with positive expression of tumor markers were annotated as malignant cells.

Copy number variation patterns in malignant cells were extracted using InferCNV(*66*) R package v3.1.5. Normal epithelial cells identified above were selected as “reference” cells for de-noise control. InferCNV analysis was performed using “denoise” mode to correct for batch effects from different mice, with tumor_subcluster_partition_method = ‘qnorm’, HMM=TRUE, and analysis_mode = ‘subclusters’. The “cluster by group” parameter was turned off to allow the observation cells to cluster unbiasedly based on CNV patterns. The CNV score for each cell was calculated as previously described(*67*). Briefly, gene expression data from the infercnv_obj@expr.data slot were scaled to the range [-1,1], and the quadratic sum of all CNV regions was computed. Malignant clusters (1–8) exhibited significantly higher CNV scores compared to the normal epithelial clusters (9–11), further validating their malignancy (fig. S3C and D).

To characterize the malignant cells, we compared gene expression of clusters 1-8 to previously identified cell identity signatures (*24, 68, 69*), embryonic stem cell-like signatures(*23*), and evolution signatures reported in a KP syngenetic transplantation model(*70*) (Fig. 2H, fig. S5). Enrichment scores of these signatures were calculated for each malignant cell using the *AUCell* algorithm implemented in SCENIC(*71*), and the scores were visualized in heatmaps. Ribosome expression was evaluated by the *AddModuleScore* function in Seurat using all ribosome genes. The number of reads per cell was calculated using SAMtools and custom Linux commands (https://kb.10xgenomics.com/hc/en-us/articles/360007068611-How-do-I-get-the-read-counts-for-each-barcode) and visualized in log2 scaled across malignant clusters. G1/S phase score was calculated by AddModuleScore function in Seurat using mouse gene set GOBP_CELL_CYCLE_G1_S_PHASE_TRANSITION from MsigDB.

RNA velocity analysis was performed to infer the dynamic states of malignant cells. Initially, spliced and unspliced transcript abundances were quantified from BAM files generated from CellRanger count using Velocyto(*72*). The resulting loom files were merged across samples. Seurat metadata was integrated with the loom file and malignant cells were extracted for downstream analysis. The dynamical model of RNA velocity in the Python module scVelo(*19*) was applied to estimate transcriptional rates and compute velocity vectors for individual cells in “stochastics” mode. The velocity fields were visualized on the batch-corrected UMAP projection, with arrows overlaid to indicate the direction and magnitude of transcriptional changes.

Trajectory analysis of malignant cells was performed using Monocle 3(*20*). Briefly, a cell data set (cds) project was constructed from the raw count matrix and cell type information of malignant cells obtained above. Batch-corrected UMAP projection was assigned to the reducedDims slot of the cds object. The trajectory was learned using the default setting in the *learn_graph* function. To determine the starting point of the trajectory, we conducted a correlation analysis between scRNA-seq epithelial malignant clusters and the three KP cell bulk RNA-seq samples. Briefly, pseudobulk expression levels of each malignant cluster were obtained using the *AggregateExpression* function in Seurat v4.0. Pearson correlation was calculated between each cluster pseudobulk vs. FPKM values of each KP bulk RNAseq sample. Cluster 2, which showed the highest correlation with KP cells, was manually selected as the root node in the graphical interface. Psedutime was measured by ordering cells according to their progress through the developmental trajectory.

### Identification of cell subtypes

We extracted T/NK cells, myeloid cells, fibroblasts, and endothelial cells for further clustering analysis, respectively. We applied resolution 1.0 for myeloid scRNA-seq data (13 clusters), 1.0 resolution for T/NK scRNA-seq data (9 clusters), 1.0 resolution for fibroblast scRNA-seq data (8 clusters), and 1.0 resolution for endothelial scRNA-seq data (3 clusters). Differential expression analysis was performed to identify cluster-specific markers as described above. Each cell cluster was annotated according to the expression profile of these markers and other canonical markers associated with different cell populations based on the literature. Markers used for annotating clusters are shown in fig. S7-S11 and described below.

Within fibroblast-lineage cell clusters (fig. S6), we first identified smooth muscle cells by *Myh11* and *Acta2*. The other fibroblast clusters were annoated by canonical markers reported in literature: pericytes (*Cspg4*, *Pdgfrb*)(*35*), adventitial fibroblasts (*Pi16*, *Mfap5*)(*35*), peribranchial fibroblasts (*Hhip*, *Aspn*)(*35*), mesothelial (*Upk3b*, *Msln*, *Wt1*)(*73*), and lipofibroblasts (*Npnt*, *Gyp*)(*73*). We also identified a cluster of cells co-expressing cancer-associated-fibroblasts markers (*Ndufa4l2*, *Postn*) and markers of a novel fibroblast subtype reported previously (*Ebf1*, *Pdzd2, Higd1b, Cox4i2*)(*73*). This cluster was annotated as *Ebf1*+ CAF. In addition, a cluster highly expressing *Cthrc1*, *Tnc*, and *Fn1* matched with Cthrc1+ fibroblasts, a subtype with high migratory capacity that was previously reported in both human and mouse lungs(*35*).

T cell clusters were first assigned to CD8+ T (*Cd8a*), CD4+T (*Cd4*), and gamma-delta T cells (*Trdc*) by corresponding canonical markers (fig. S9). Then the CD8+ T clusters were further annotated to subtypes based on the expression of markers previously with different CD8+ T cells states and function(*74, 75*): naïve (*Tcf7*, *Ccr7*, *Il7r*, *Sell*), effector-memory (*Eomes*, *Gzmk*, *Ccl5*, *Ccr5*), interferon-stimulated genes (Isg)-positive (*Isg15*, *Isg20*, *Ifit3*, *Ifit1*), and kilter cell immunoglobulin-like receptor (Kir)-positive (*Klrc2*, *Klra7*, *Klrd1*). Similarly, CD4+ T clusters were annotated as follows: naïve (*Tcf7*, *Ccr7*, *Il7r*, *Sell*), follicular helper (Tfh) (*Tnfrsf11, Tnfrsf8, Slamf6*), *Ifng*+ T helper 1 (Th1) (*Ifng*, *Havcr2*, *Lag3*, *Gzmb*), and regulatory T (Treg) (*Foxp3*, *Ctla4*, *Il2ra*).

Within myeloid clusters (fig. S10), we first identified cDC1 (*Xcl1*, *Cd36*, *Itgae*), cDC2 (*Itgax*, *H2-DMb1*, *Mgl2*), mature regulatory DC (mregDC) (*Fscn1*, *Ccl22*, *Cacnb3*), plasmacytoid DC (pDC) (*Siglech*, *Ccr9*, *Bsl2*), 2 clusters of neutrophils (*S100a8*, *S100a9*, *Cxcl2*), 2 clusters of monocytes (*Lyz2*, *Vcan*, *Chil3*, *Fn1*), and 5 clusters of macrophages (*Cd68*, *Mrc1*, *C1qc*, *C1qb*). The neutrophil clusters were further annotated as *Retnlg*+ and *Cxcl2*+ as previously described(*76*), where *Retnlg*+ neutrophils were characterized by high expression of canonical neutrophil marks (*Csf3r*, *S100a8*, *G0s2*), while *Cxcl2*+ neutrophils expressed *Ccl3*, *Cclr2* and were reported related to homing of tumor-associated neutrophils(*77*) and neutrophils aging(*78*). Monocytes clusters were further named as Ly6c2+ and Ly6c2-subtypes(*76*). The macrophages were fully annotated as follows: first, the 5 clusters were identified as alveolar macrophages (AM) (*Atp6v0d2, Plet1, Lpl, Ctsd*) and interstitial macrophages (IM) (*C1qa, C1qb, C1qc*) by their markers(*79*); second, a subtype of AM cells expressing lipid-associated genes (*Gpnmb*, *Ctsb*, *Ctsl*, *Lgals3*)(*79*) was defined as Lipid-associated alveolar, and the other AM cluster was found matching with alveolar resident cells which highly expressed *Krt79*, *Krt19*, *Car4*, and *Chil3*(*79*); third, interstitial subtypes were confirmed by specific marker genes including inflammatory cytokine-enriched (*Ccr5*, *Il1b, Nrkb1*), pro-angiogenic (*Fn1, Thbs1, Arg1*), and a proliferative cluster featured by ribosome genes and enriched for MYC-target gene set.

Subtypes of endothelial cells were identified using the following markers(*80*): capillary A (*Sema3c, Gpihbp1,Pttp, Plvap*), capillary B (Igfbp7, Car4, Emp2), and blood vessel endothelial (*Pf4*, *Nrgn*, *Alox12*) (fig. S11).

### Stemness and ADI scores

A stemness score was established via *AddModuleScore* in Seurat using the genes from the pathways “Mechanisms associated with Pluripotency” (WP), “Cell fate specification” (GO) and “cell fate commitment” (GO) obtained via GSEA analysis of malignant cells from young versus old KP mice in Table S5. The ADI signature score was composed of 10 established marker genes from the literature(*46*) as well as non-overlapping marker genes from our own dataset within the top 10 markers of the epithelial proliferative population (Table S8). We then assessed the expression of the genes included in the ADI score in scRNA-Seq data from 61 human NSCLC samples(*47*), and sorted them based on age. Briefly, enrichment of ADI signatures was calculated by *AUCell* for each epithelial cell extracted from the NSCLC samples. Then patient-level ADI score was summarized as the mean across epithelial cells. A Chi^2^-test was employed to determine if a high ADI score was associated with advanced age. Age of NSCLC human samples(*47*) was divided into two groups based on the median age (64). ADI scores were binned into three equal quantiles, whereby low ADI score<0.04, medium ADI score=0.04-0.14, and highADI score ≥0.14.

### Visualization of gene signatures

Marker genes were visualized on UMAP projections or violin plots using log-normalized counts. For the bubble plot of marker genes, the average expression of each gene was calculated for each cluster/group and then normalized by mean and standard deviation (z-scores) and the size of bubble was used to denote percentage of cells expressing the markers in cluster/group. To systematically assess the effects of aging, differential expression analysis was performed comparing young vs. old cells within each major cell population, as well as within subtypes of T/NK cells, myeloid cells, fibroblasts, endothelial cells, and epithelial cells. Following differential analysis, pre-ranked gene set enrichment analysis (GSEA) was performed. Genes were ranked based on -log10(p-value)*(sign of log2(fold-change)), positioning the most upregulated genes at the top and the most downregulated genes at the bottom of the ranking. Pre-ranked GSEA was performed on gene rankings using R package fgsea (*81*) with 10,000 permutations against Hallmark, REACTOME, and GO databases from MsigDB (*82–84*). The normalized enrichment scores (NES) of gene sets were visualized using heatmaps.

### Analysis and visualization of cell-cell interaction using “CellChat”

Cell-cell communication analysis was preformed using CellChat (*39*) v2. Briefly, scRNA-seq data were first loaded into the CellChat R package with two biological conditions (young vs. old). The database CellChatDB.mouse from CellChatDB (*39*) v2 was used as the ligand-receptor interaction database for mouse data. The young dataset and the old dataset were processed separately in the following steps: scRNA-seq expression data were preprocessed using functions *identifyOverExpressedGenes* and *identifyOverExpressedInteractions*; the communication probability between each cell type within the same biological condition was computed and filtered using functions *computeCommunProb* with the method “trimean”, and *filterCommunication* with min.cells=9; the pathway-level communication probability was computed using function *computeCommunProbPathway*; the aggregated cell-cell communication network was calculated by function *aggregateNet*; the network centrality scores were computed by the function netAnalysis_*computeCentrality*. The young and old datasets were then merged into one CellChat object for further comparison between different biological conditions. The number of interactions and interaction strengths between each cell type were compared among young conditions and old conditions. The signaling network similarity for the young condition and old condition was computed using the function *computeNetSimilarityPairwise*. Manifold learning and classification learning of the signaling networks were conducted by *netEmbedding* and *netClustering*, respectively. Cell-cell communication analysis results were visualized using CellChat. The average interaction strength from CellChat was used as the cell-cell interaction score between different cell groups. Interactions between the sender and the receiver were calculated separately.

### Statistical Analysis

Statistical analyses for mouse and histology experiments were performed using GraphPad Prism 9 (GraphPad Software, Boston MA, USA). Comparisons between the two groups were made using unpaired Student’s T-tests. Statistical significance was defined as p<0.05. For datasets with potential outliers, we employed the ROUT method (Q=1%) to objectively identify outliers using GraphPad Prism. Data points were excluded if they were confirmed as outliers by this test.

## Data Availability

Bulk RNA-Seq data presented in this study have been deposited under GEO accession code GSE286493. Single-cell RNA-Seq data presented in this study have been deposited under GEO accession code GSE286494. All other data can be made available upon reasonable request to.

## Acknowledgements

We thank members of the Gomes lab and Dr. Gina DeNicola for their helpful feedback and Drs. Alfred Zippelius (University of Basel) and Massimo Broggini (Instituto di Ricerche Farmacologiche Mario Negri IRCCS) for kindly sharing KP1.9 cells. We thank Brooke Jackson (Moffitt) for helping with immunofluorescence analysis and Sean Yoder (Moffitt) for help with planning single-cell RNA-Sequencing experiments. We would also like to thank the Moffitt Cancer Center/USF Comparative Medicine Program for animal care and staff members of the Flow Cytometry, Analytical Microscopy, Molecular Genomics core facilities at the H. Lee Moffitt Cancer Center & Research Institute, an NCI designated Comprehensive Cancer Center (P30-CA076292). This work was directly supported by a New Innovator Award from OD/NIH to APG (DP2AG0776980). APG and the Gomes Laboratory are also supported by an American Cancer Society Research Scholar Award (RSG-22-164-01-MM), the NIA (R21AG083720), the NCI (R01CA279023), the Florida Health Department Bankhead-Coley Research Program (24B03), METAvivor, the Florida Breast Cancer Research Foundation, and the Phi Beta Psi Sorority. SD was supported by a Miles for Moffitt Postdoctoral Fellowship and is currently supported by a Postdoctoral Fellowship from the American Cancer Society (PF-24-1151991-01-MM). Schematic figures were created using BioRender.com.

## Ethics Declaration

The authors have no competing interests to disclose.

---

**Figure S1 Lung cell type composition flow cytometry gating strategy**

**A.** Flow cytometry gating strategy for immune cells from young and old lungs. **B.** Flow cytometry gating strategy for non-immune cells from young and old lungs.

**Figure S2 Optimization of a KP syngeneic transplantation model**

**A.** Representative hematoxylin & eosin (H&E) staining images for grade 3, 4 and 5 tumors in our syngenetic KP transplantation model versus the autochthonous GEMM model. **B.** UMAP projection of malignant cells from young transplanted mice, colored by cluster. **C.** Heatmap showing similarity of clusters from KP syngeneic transplantation model compared to clusters from the autochthonous GEMM model (Marjanovic et al. *Cancer Cell*, 2020) **D.** Representative H&E images of young and old mouse lungs 4 weeks after KP cell transplantation. Yellow outlines represent total lung area annotations, and red outlines represent tumor annotations. **E.** Quantification of tumor burden in transplanted KP tumor-bearing lungs from young and old mice (n=20 young and 16 old mice, t-test). **F.** Quantification of tumor number in transplanted KP tumor-bearing lungs from young and old mice (n=20 young and 16 old mice, t-test). **G.** Representative hematoxylin & eosin (H&E) staining images for grade 3, 4 and 5 tumors in young and old mice transplanted with KP cells. **H.** Quantification of tumor grade in tumors from young and old mice transplanted with KP cells (n=19 young and 16 old mice, t-test). Data are presented as the mean ± SD.

**Figure S3 scRNA-Seq quality control**

**A.** UMAP projection of all TME cells from tumors isolated from young and old KP tumor-bearing mice, colored by mouse biological replicate **B.** Bubble plot comparison of KP transplant clusters versus Tumor, KP, Cell Cycle, and Atlas of the Aging lung marker genes **C.** Copy Number Variation (CNV) score in each cluster of epithelial malignant cells **D.** Estimation of copy number variants by InferCNV of KP epithelial malignant cells (bottom) with normal epithelial cells were used as control (top) **E.** Heatmap showing z-scores for the expression of various ribosome genes across clusters of epithelial malignant cells **F.** Heatmap showing the expression of common housekeeping genes across the different clusters of epithelial malignant cells

**Figure S4 Transplantation Induces Different Subpopulations with Varying Similarities to Parental KP Cells and Differential Expression of Key KP-Related Genes**

**A.** Pearson Correlation between pseudobulk gene expression of each cluster of epithelial malignant cells and the bulk gene expression of cultured parental KP cells. **B.** UMAP projection showing gene expression for select genes involved in epithelial lineage and stemness.

**Figure S5 Comparison of epithelial malignant cells from KP transplant mice with cell identity signatures**

**A.** Heatmap of transcriptional similarity between scRNA-Seq clusters of epithelial malignant cells isolated from young and old mice and embryonic gene signatures(*23*) **B.** Heatmap of transcriptional similarity between scRNA-Seq clusters of epithelial malignant cells isolated from young and old mice and gene signatures of different cell types(*24*) **C.** Heatmap of transcriptional similarity between scRNA-Seq clusters of epithelial malignant cells isolated from young and old mice and gene signatures of different trajectories involved in organogenesis(*24*) **D.** Heatmap of transcriptional similarity between scRNA-Seq clusters of epithelial malignant cells isolated from young and old mice and gene signatures of difference cell types(*86*). Shown are z-score enrichment scores.

**Figure S6 Aging drives LUAD’s gain of stemness characteristics**

**A.** Violin Plots showing the expression of stemness markers by scRNA-Seq subcluster of malignant cells **B.** Violin Plots showing number of reads per cell, number of gene per cell, Log10(Genes per UMI), G1, G2M, S, and ribosome score across different malignant clusters **C.** Representative IHC images of young and old KP tumor-bearing lungs stained for RNA-Pol II p-Ser5 (left), with associated thresholding mask (right). **D.** Quantification of RNA-Pol II p-Ser5^High^ cells in tumors from young and old KP tumor-bearing lungs (n=31 tumors from 3 young mice, n=53 tumors from 5 old mice, t-test). Dotted lines in violin plots represent the median and quartiles.

**Figure S7 ScRNA-Seq cluster identification of TME cells from mice transplanted with KP cells**

**A.** UMAP projection of TME cells showing the expression of marker genes for each major cell type **B.** Bubble plot showing marker genes used to identify each cell type **C.** Violin plots showing number of unique genes detected, number of reads and S-phase score for each scRNA-Seq sample, stratified by mouse biological replicate **D.** UMAP projection of TME cells colored by cluster and separated by age **E.** Pie charts showing the proportions of major cell types within the TME of young vs old mice.

**Figure S8 Subcluster identification of fibroblasts derived from KP transplanted mice**

**A.** UMAP projection of fibroblast cells showing the expression of marker genes for various subtypes **B.** Bubble plot showing marker genes used to identify each fibroblast cell subcluster **C.** Violin plots showing number of unique genes detected, number of reads and S-phase score for each fibroblast subtype **D.** UMAP projection of fibroblast cells from young and old KP tumors, colored by subcluster **E.** Bar plot showing proportions of fibroblast subclusters in young and old mouse KP tumors **F.** Quantification of Tnc^+^/Pdgfrα^+^ cells in young versus old KP-tumor bearing mice (n=39 tumors from 5 young mice, n=55 tumors from 5 old mice, t-test). Dotted lines in violin plots represent the median and quartiles. **G.** Quantification of the percentage of Tnc^+^/Pdgfrα^+^ cells over all Pdgfrα^+^ fibroblasts in young versus old KP-tumor bearing mice (n=39 tumors from 5 young mice, n=55 tumors from 5 old mice, t-test). Dotted lines in violin plots represent the median and quartiles. **H.** Representative immunofluorescence images of Tnc, Pdgfrα and DAPI staining of lung samples from young and old KP tumor-bearing mice. Dotted squares represent the area shown with staining deconvoluted into different channels.

**Figure S9 Subcluster identification of T cells derived from KP transplanted mice**

**A.** UMAP projection of T cells showing the expression of marker genes CD4, CD8 and gd T cells **B.** Bubble plot showing marker genes used to identify each CD4 T cell subcluster **C.** Bubble plot showing marker genes used to identify each CD8 T cell subcluster **D.** Violin plots showing number of unique genes detected, number of reads and S-phase score for each T cell subtype **E.** UMAP projection of T cells colored by subtype cluster **F.** Bar plot showing proportions of T cell subclusters in young and old mouse KP tumors **G.** Quantification of the number of NKG2A^+^/CD8^+^ cells over area in young versus old KP-tumor bearing mice (n=15 tumors from n=3 young mice, n=36 tumors from n=5 old mice, t-test) **H.** Quantification of the percentage of NKG2A^+^/CD8^+^ cells over all CD8^+^ T cells in young versus old KP-tumor bearing mice (n=17 tumors from n=3 young mice, n=37 tumors from n=5 old mice, t-test). Dotted lines in violin plots represent the median and quartiles. **I.** Representative immunofluorescence images of CD8, NKG2A and DAPI staining of lung samples from young and old KP tumor-bearing mice. Dotted squares represent the area shown with staining deconvoluted into different channels.

**Figure S10 Subcluster identification of myeloid cells derived from KP transplanted mice.**

**A.** UMAP projection of myeloid cells colored by subtype cluster **B.** UMAP projection of myeloid cells showing the expression of marker genes for various myeloid subtypes **C.** Bubble plot showing marker genes used to identify each macrophage subcluster **D.** Bubble plot showing marker genes used to identify each myeloid cell subcluster **E.** Violin plots showing number of unique genes detected, number of reads and S-phase score for each myeloid subtype **F.** Bar plot showing proportions of myeloid subclusters in young and old mouse KP tumors.

**Figure S11 Subcluster identification of endothelial cells derived from KP transplanted mice**

**A.** UMAP projection of endothelial cells colored by subtype **B.** UMAP projection of endothelial cells showing the expression of marker genes for various endothelial cell subtypes **C.** Bubble plot showing marker genes used to identify each endothelial subcluster **D.** Violin plots showing number of unique genes detected, number of reads and S-phase score for each endothelial subtype **E.** Bar plot showing proportions of epithelial subclusters in young and old mouse KP tumors.

**Figure S12** – **Proliferative epithelial cells are the most communicative cell type within the aged TME**

**A.** UMAP projection of normal epithelial cells, colored by subcluster. **B**. Violin plot showing the gene expression level of *Sftpc* in malignant compared to normal epithelial cells **C.** Heatmap of CellChat interactions strength between the different cell types within the KP TME **D.** Chord diagrams showing interaction number and strength between of different cell types within the TME, with epithelial normal cells deconvoluted **E.** Chord diagrams showing number and strength of interactions originating from other cell types onto the epithelial proliferative cells. Circle size represents the number of each cell type detected **F.** Chord diagrams showing number and strength of interactions originating from epithelial proliferative cells onto other cell types within the TME. Circle size represents the number of each cell type detected.

**Figure S13 Aging leads to an increase in Alveolar Differentiation Intermediate cells upon KP transplantation.**

**A.** Quantification of AT2 cells in tumor-adjacent areas (200µm radius around tumor perimeter) from young and old KP tumor-bearing mice, defined as SPC^+^/P53^+^ or SPC^+^/Pdpn^-^/Krt8^-^ cells (n=42 tumors from 5 young mice, n=48 tumors from 5 old mice, t-test) **B.** Quantification of AT1 cells in tumor-adjacent areas (200µm radius around tumor perimeter) from young and old KP tumor-bearing mice, defined as Pdpn^+^/P53^+^ or Pdpn^+^/Spc^-^/Krt8^-^ cells (n=42 tumors from 5 young mice, n=48 tumors from 5 old mice, t-test) **C.** Quantification of the number of Krt8^+^/P53^+^ cells in young versus old non-tumor-bearing (NTB) mouse lungs **D.** Chord Diagram from CellChat data showing signaling interactions originating from ADI cells towards epithelial malignant cells. **E.** Pie chart showing the categories of cell-cell interactions originating from ADIs towards the malignant cells (related to Table S9) **F.** Violin plots showing the log2(Normalized gene expression) for Wnt5a and Jag1 in epithelial subtypes from KP tumor-bearing mice.

## Supplementary Materials

**Table S1.** Gene Set enrichment analysis of RNA-Seq data from non-tumor bearing lungs from 3-month-old versus 24-month-old mice from Kawaguchi K, et al. *Exp Anim* (2023)

**Table S2.** List of marker genes for each cluster of epithelial malignant cells isolated from young and old KP tumor-bearing mice.

**Table S3.** List of differentially expressed genes in malignant epithelial cells isolated from young versus old KP tumor-bearing mice.

**Table S4.** Table of the number of unique molecular indices (UMIs) per gene detected via single-cell RNA-Sequencing of malignant epithelial cells isolated from young and old KP tumor-bearing mice.

**Table S5.** Gene Set enrichment analysis of epithelial malignant cell isolated from young versus old KP-tumor bearing lungs.

**Table S6.** List of differentially expressed genes between young and old KP tumor-bearing mice for all major cell types within the TME

**Table S7.** List of differentially expressed genes between young and old KP tumor-bearing mice for all minor cell types within the TME

**Table S8.** List of marker genes for each subpopulation of normal epithelial cells.

**Table S9.** List of CellChat interactions between ADIs/epithelial proliferative cells and other cells within the TME

**Table S10.** Gene Set enrichment analysis of genes differentially expressed in ADIs compared to other epithelial normal cells.

**Table S11.** List of genes constituting the ADI signature score

**Table S12.** Gene Set enrichment analysis of secreted factors derived from ADIs.

