## Supplementary figures and images for "Aging directs the differential evolution of KRAS-driven lung adenocarcinoma"

### fig. S1

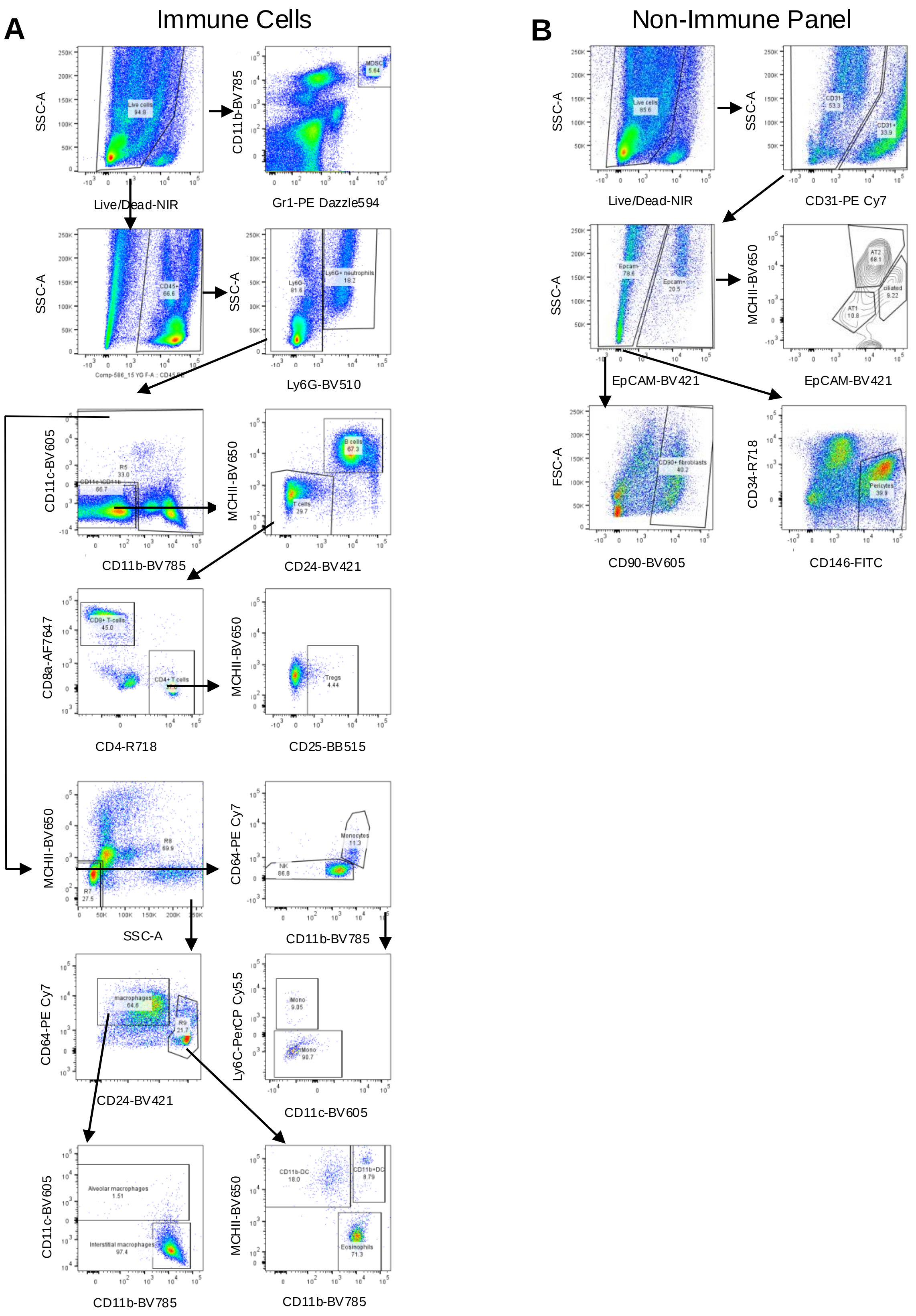

### fig. S2

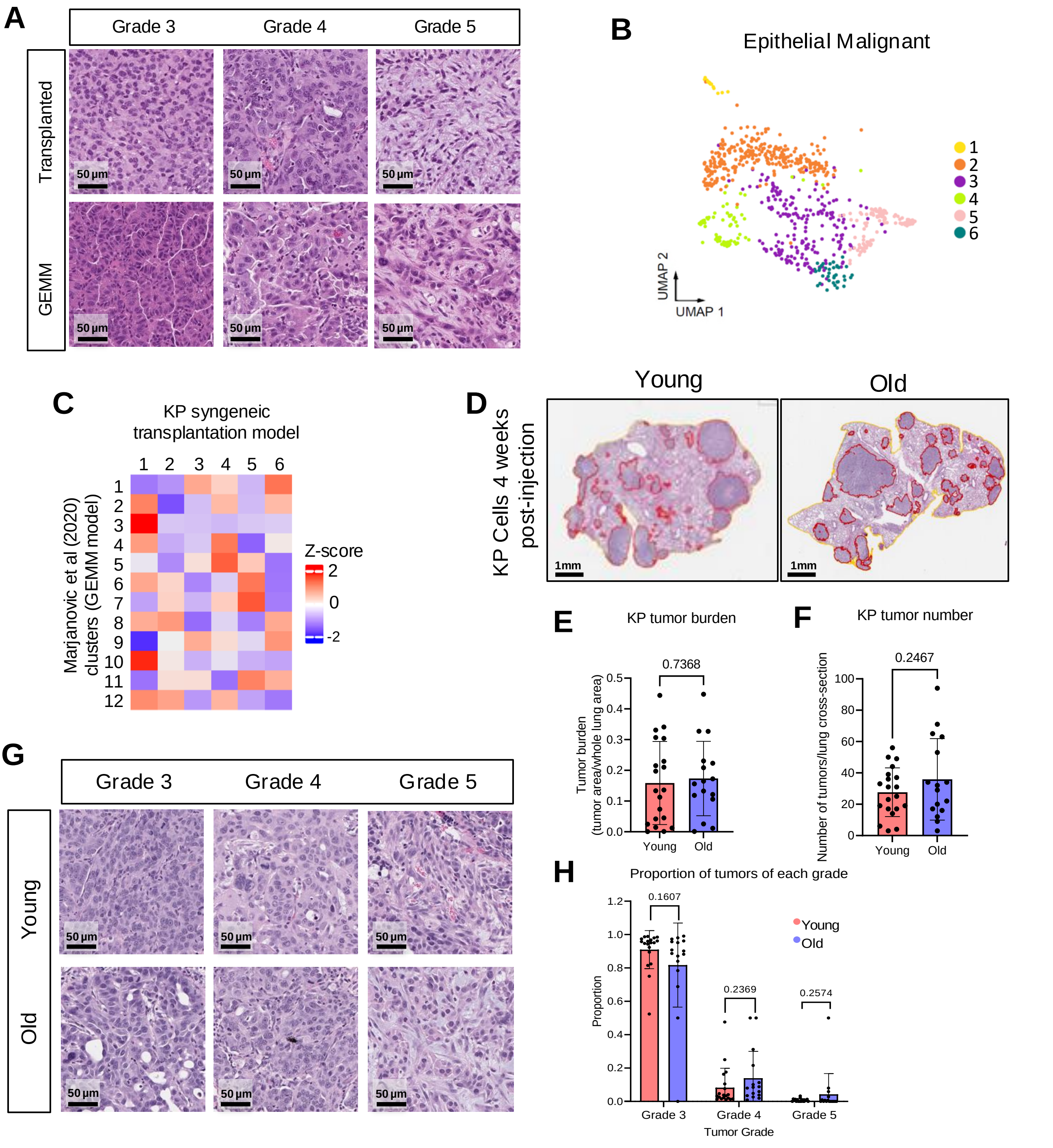

### fig. S3

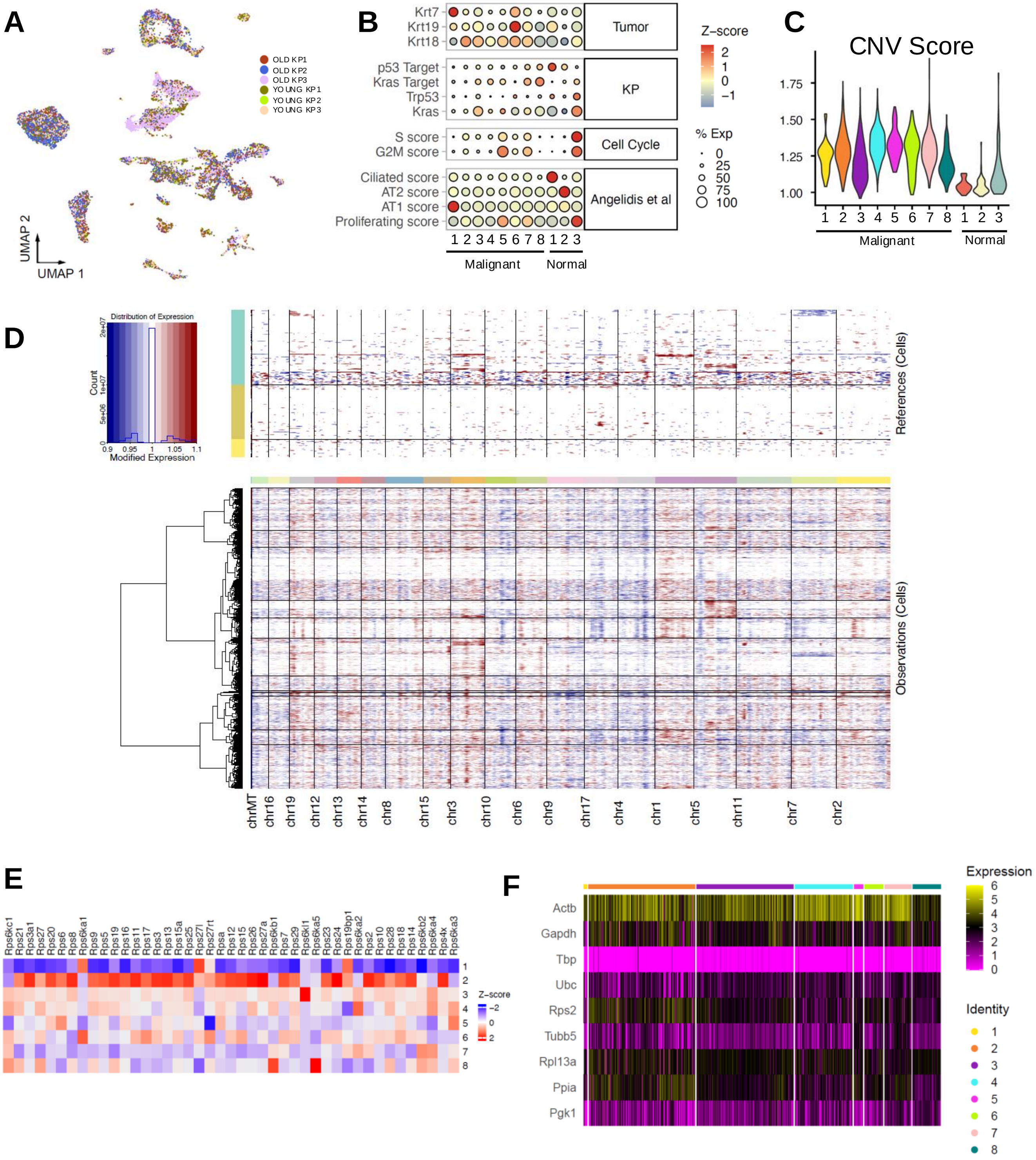

### fig. S4

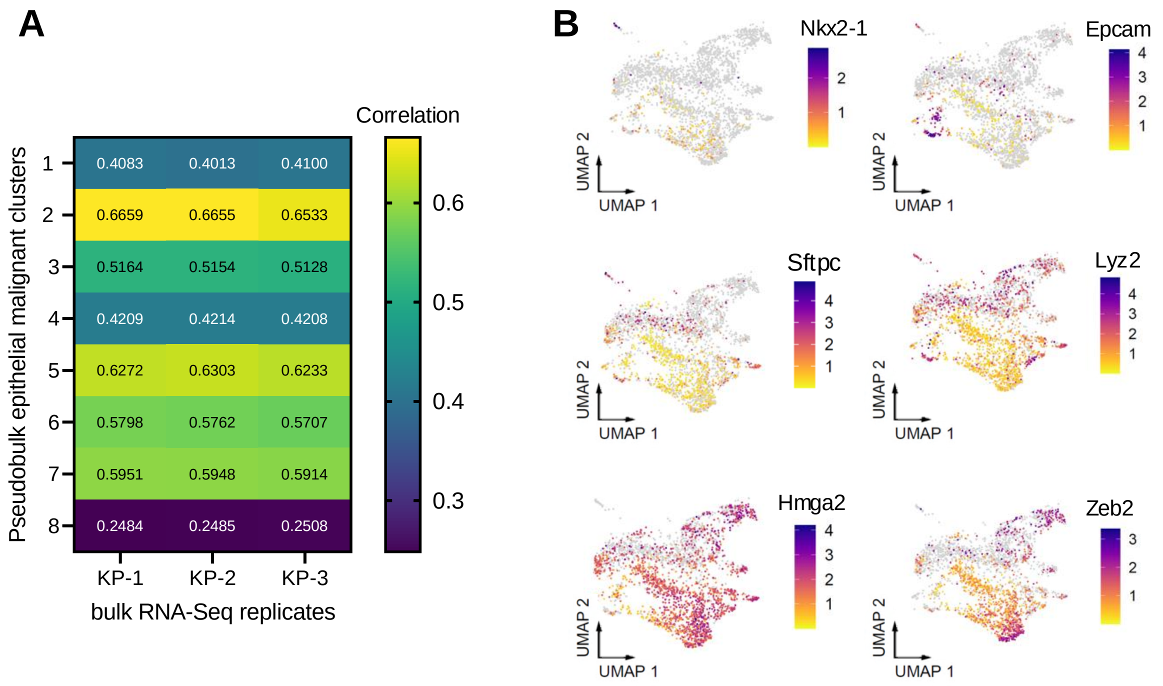

### fig. S5

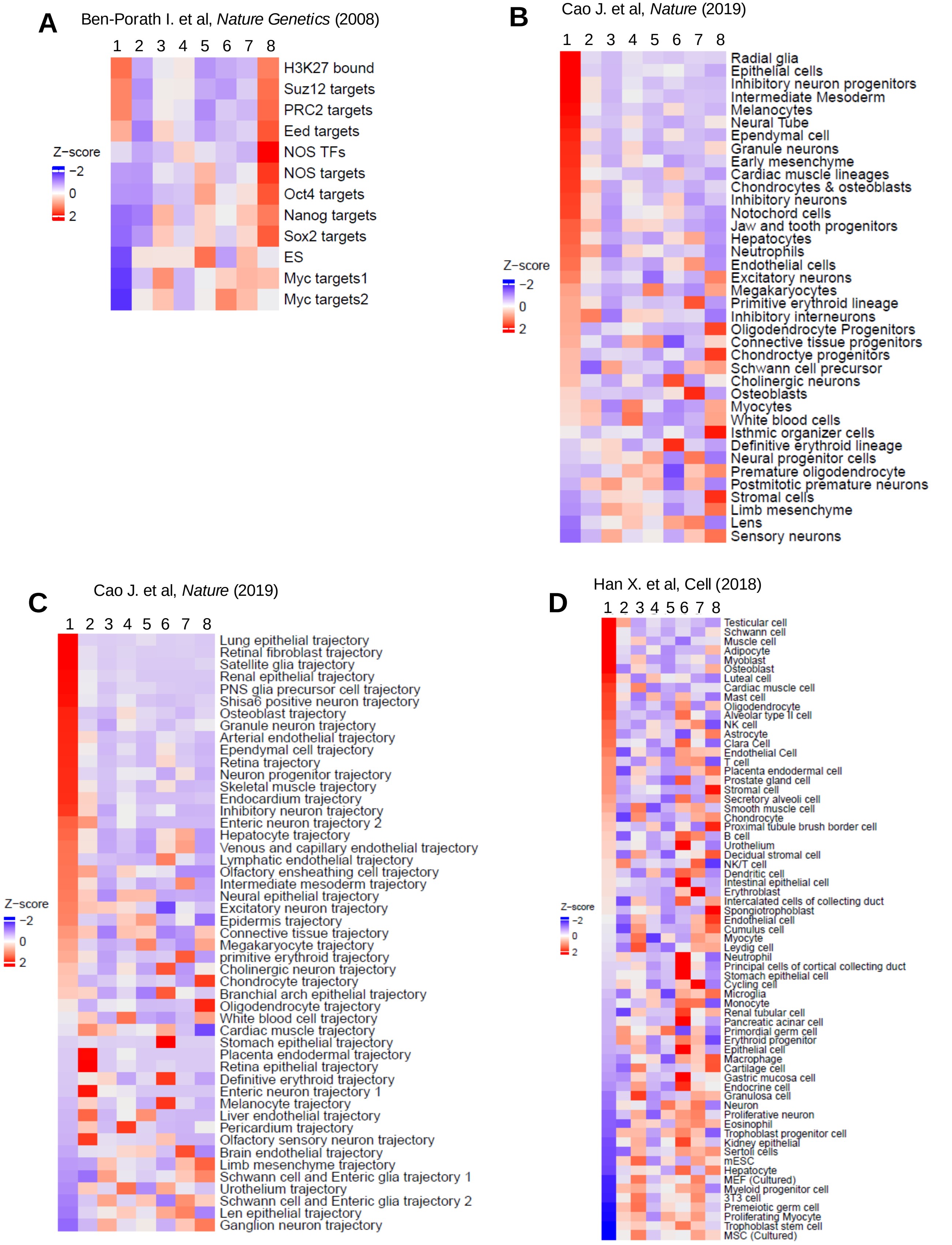

### fig. S6

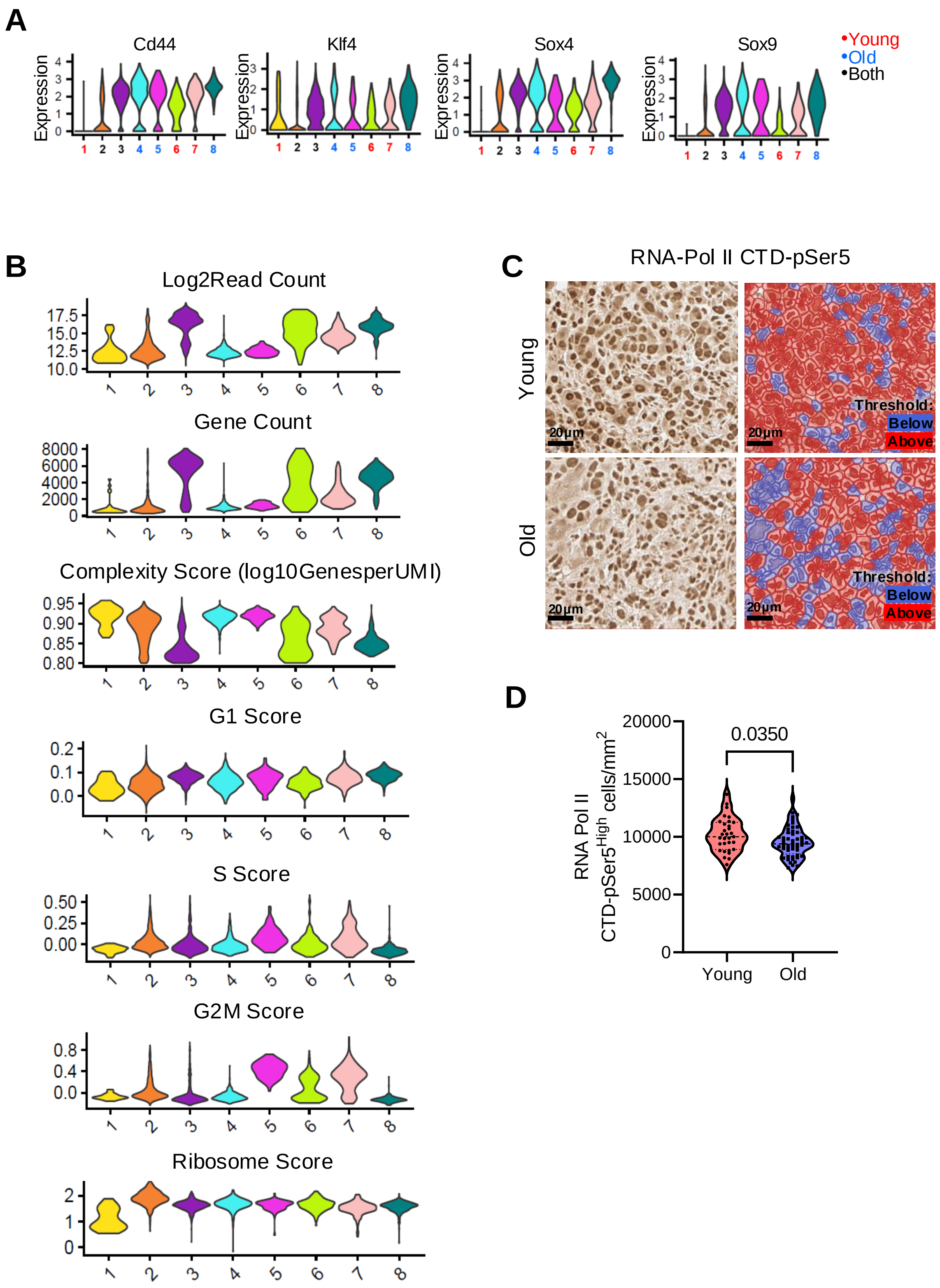

### fig. S7

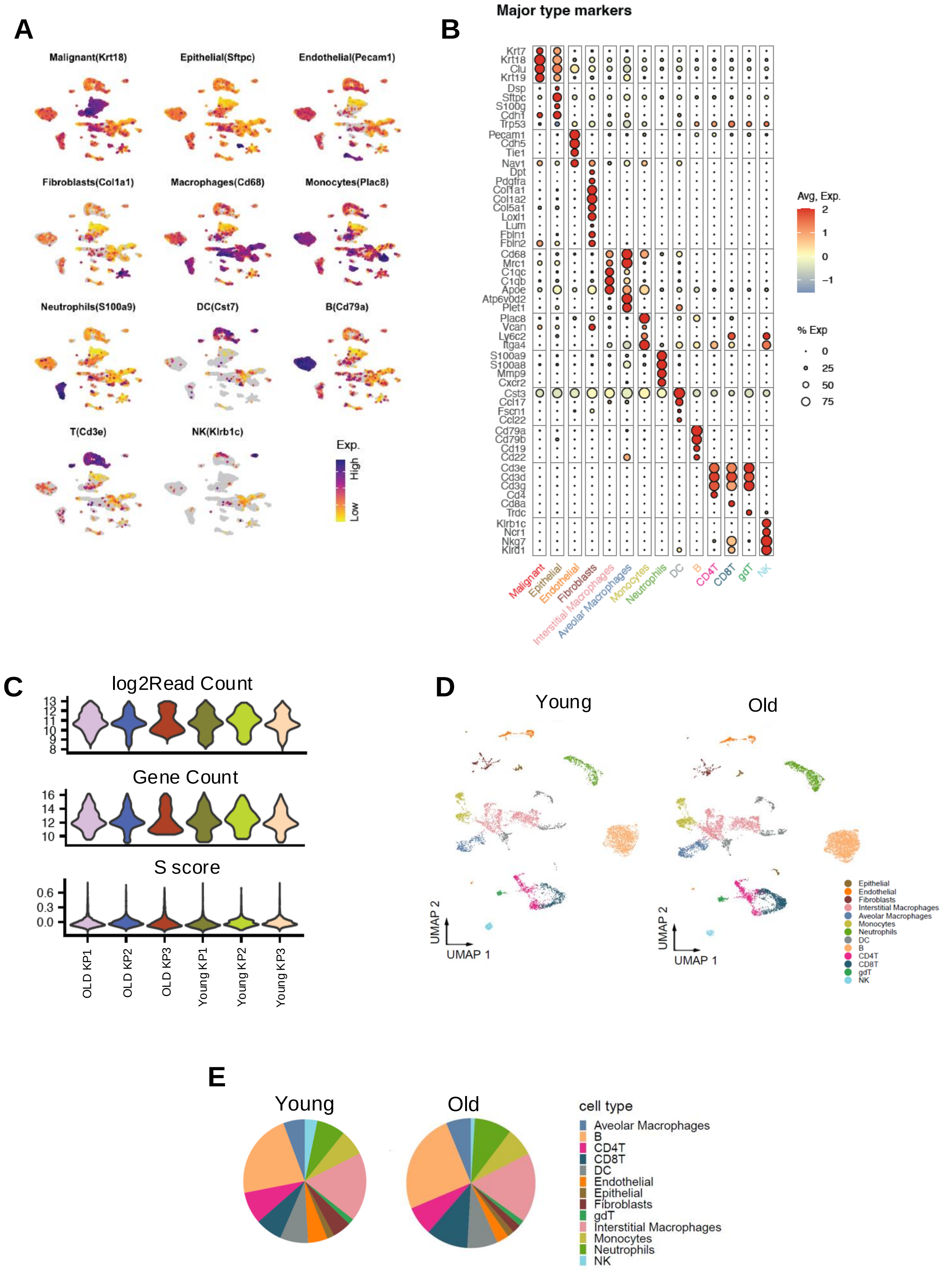

### fig. S8

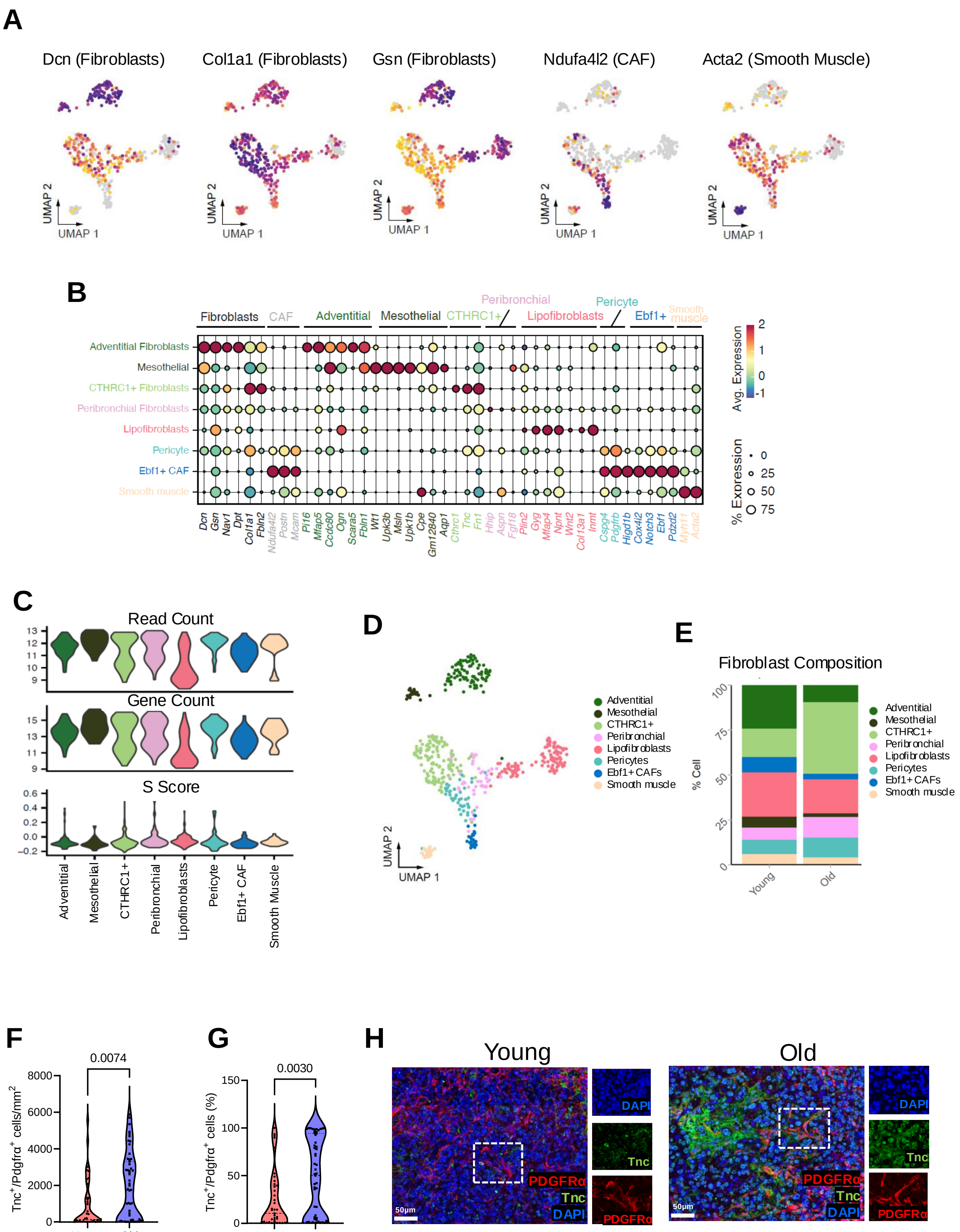

### fig. S9

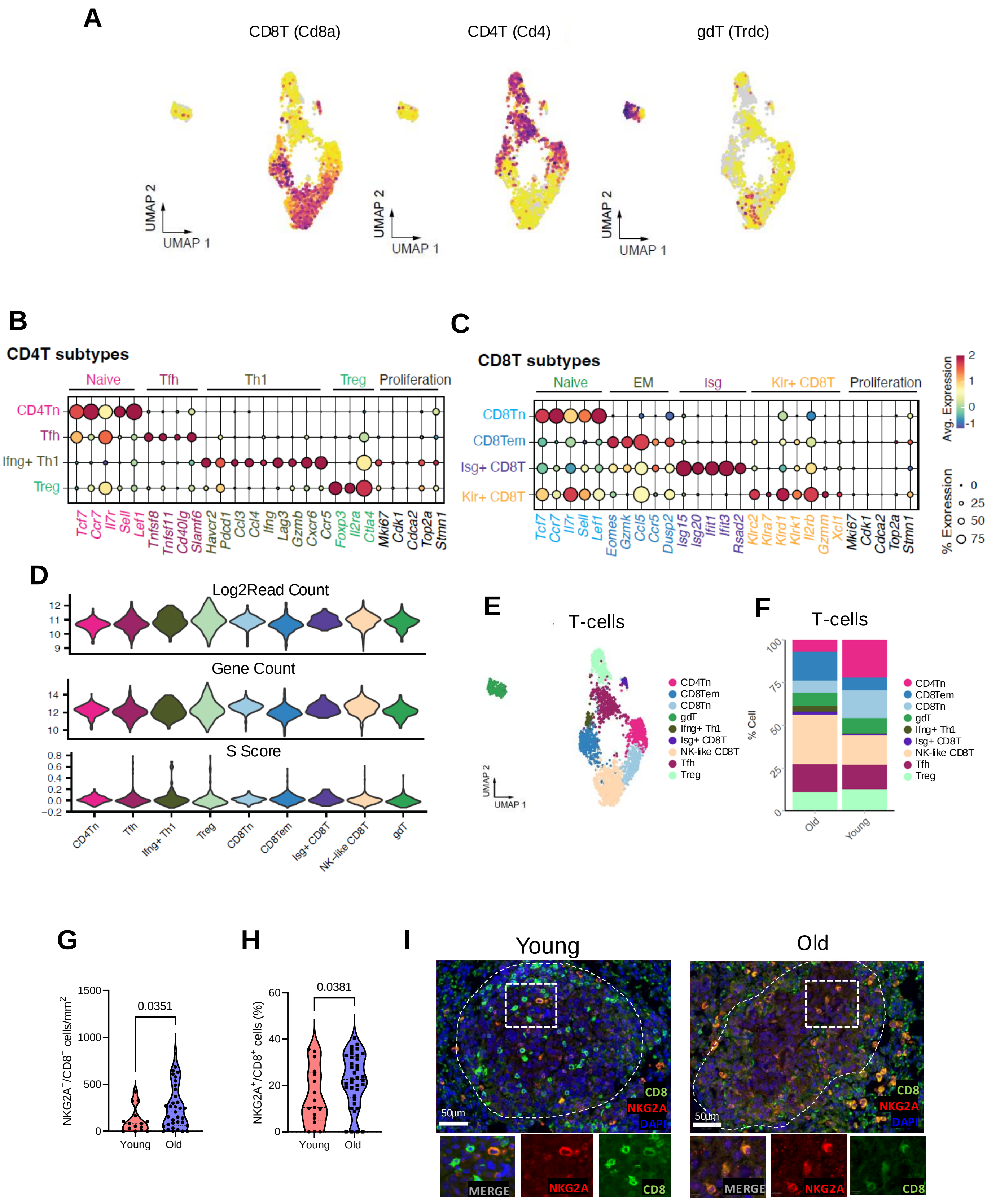

### fig. S10

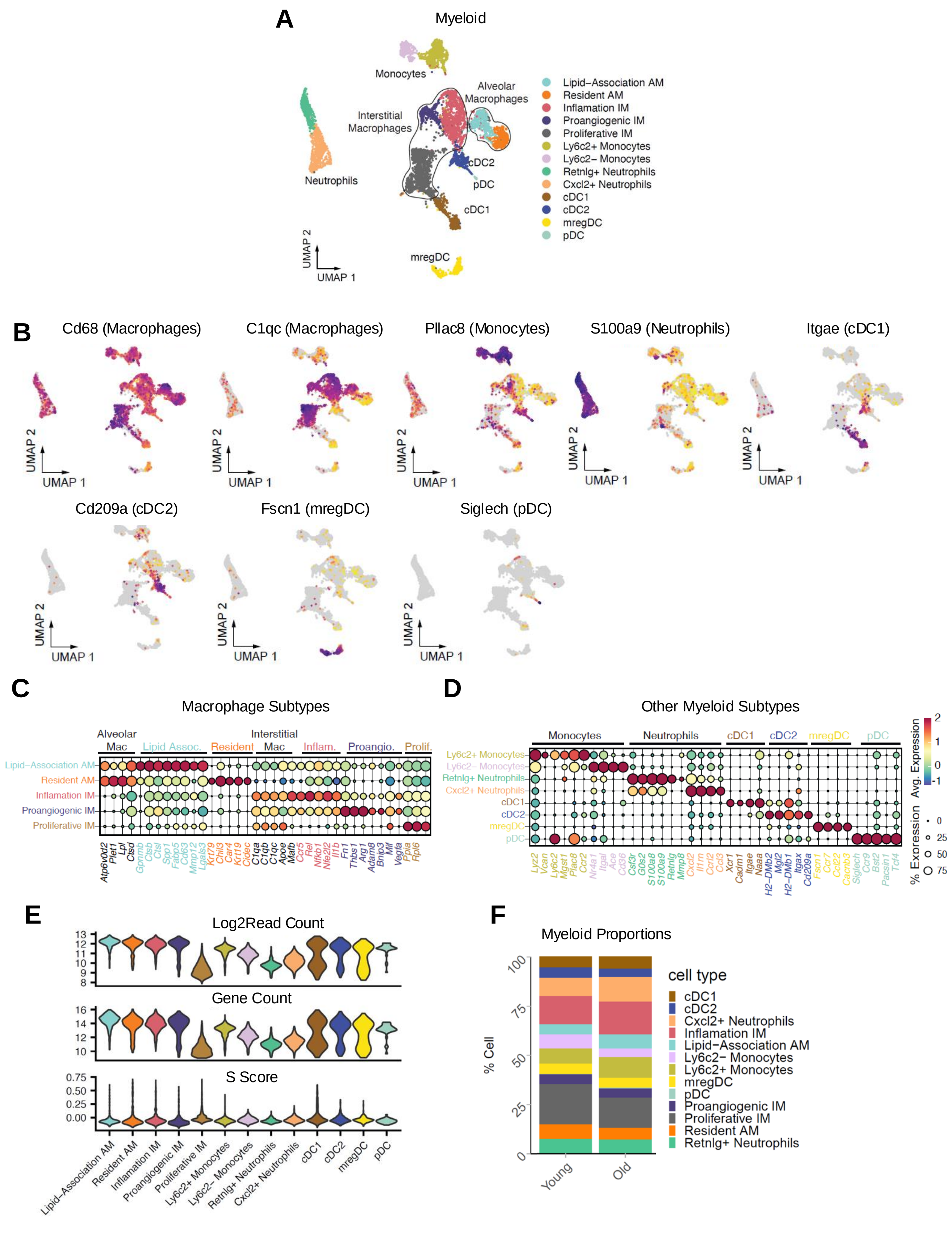

### fig. S11

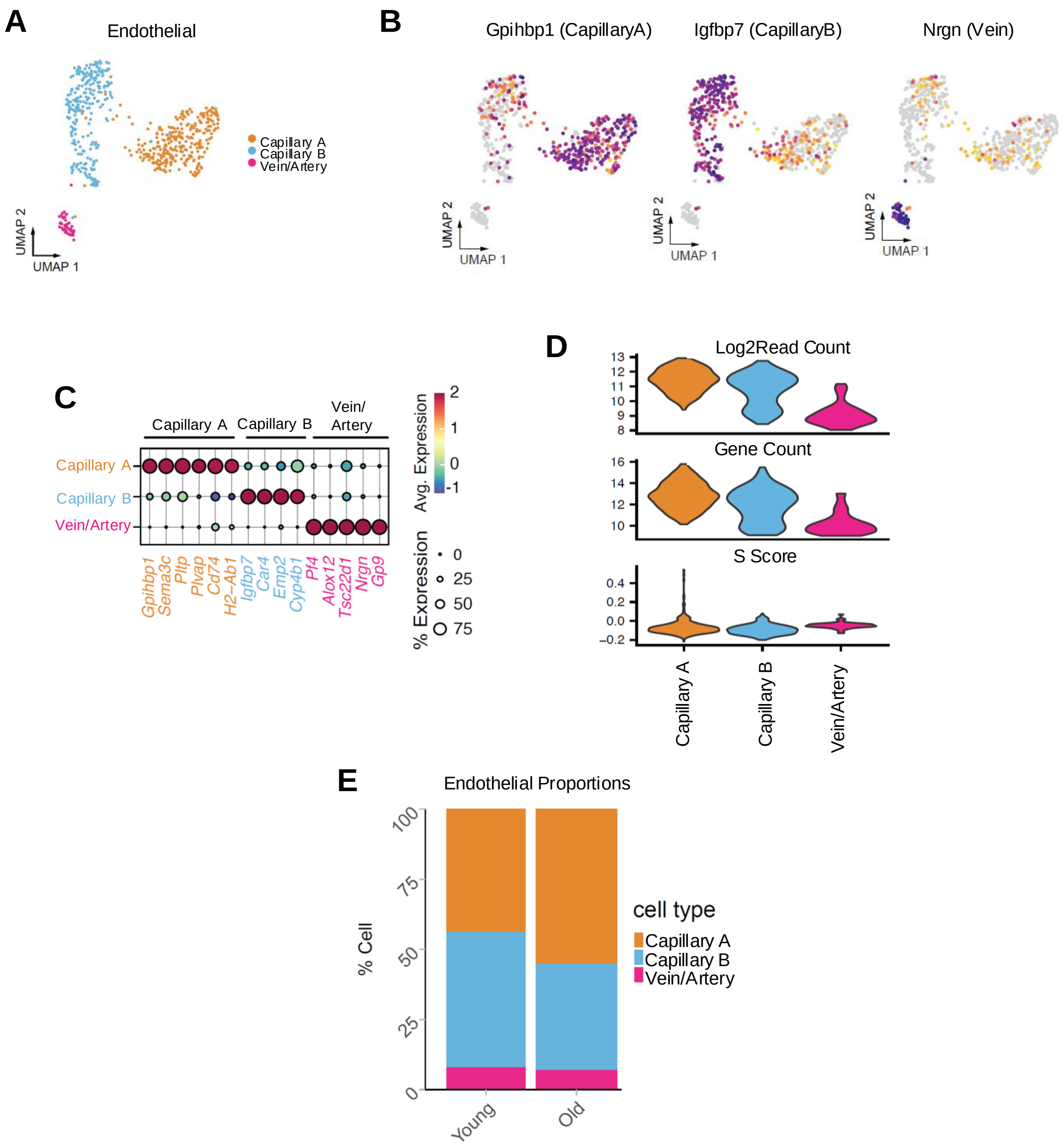

### fig. S12

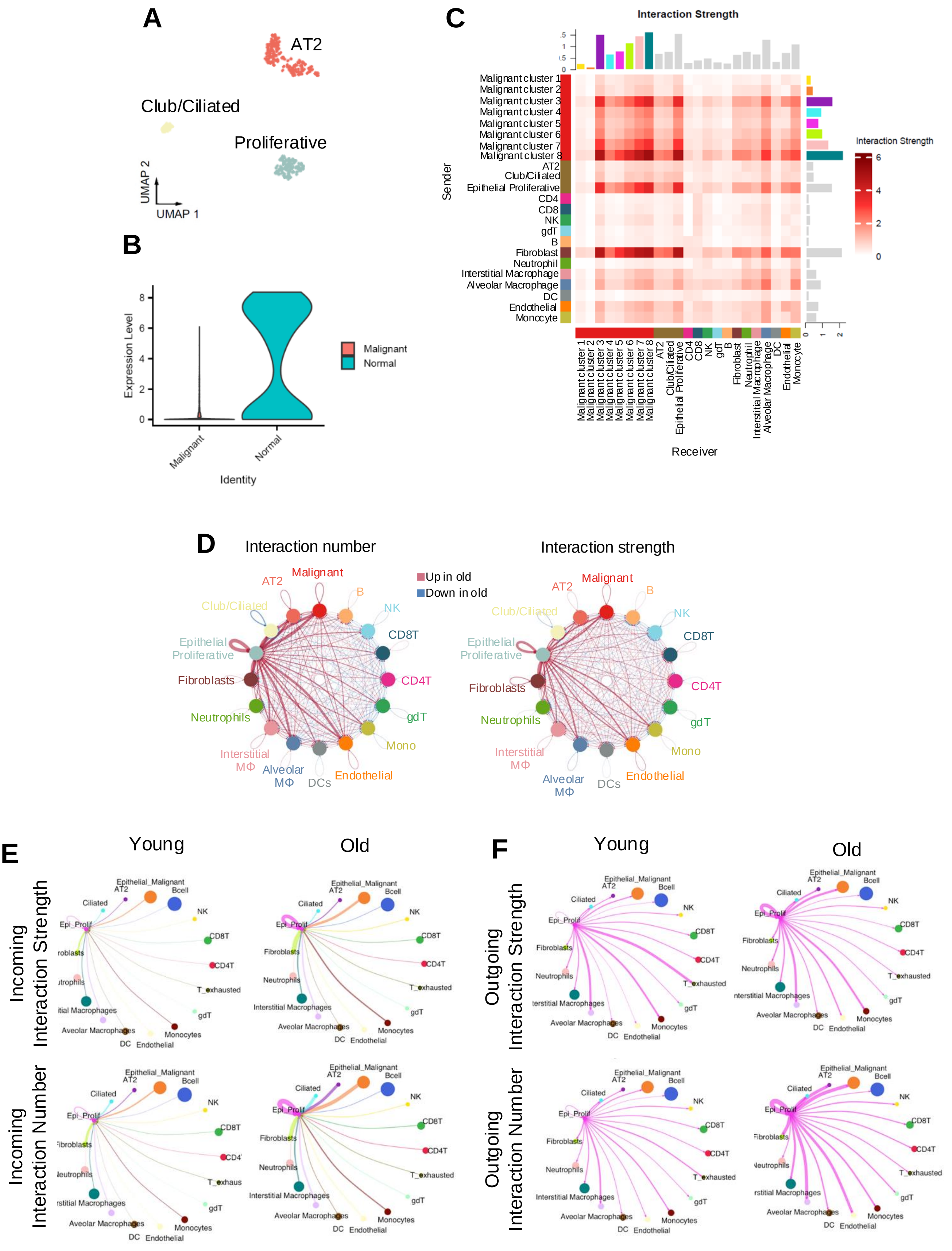

### fig. S13

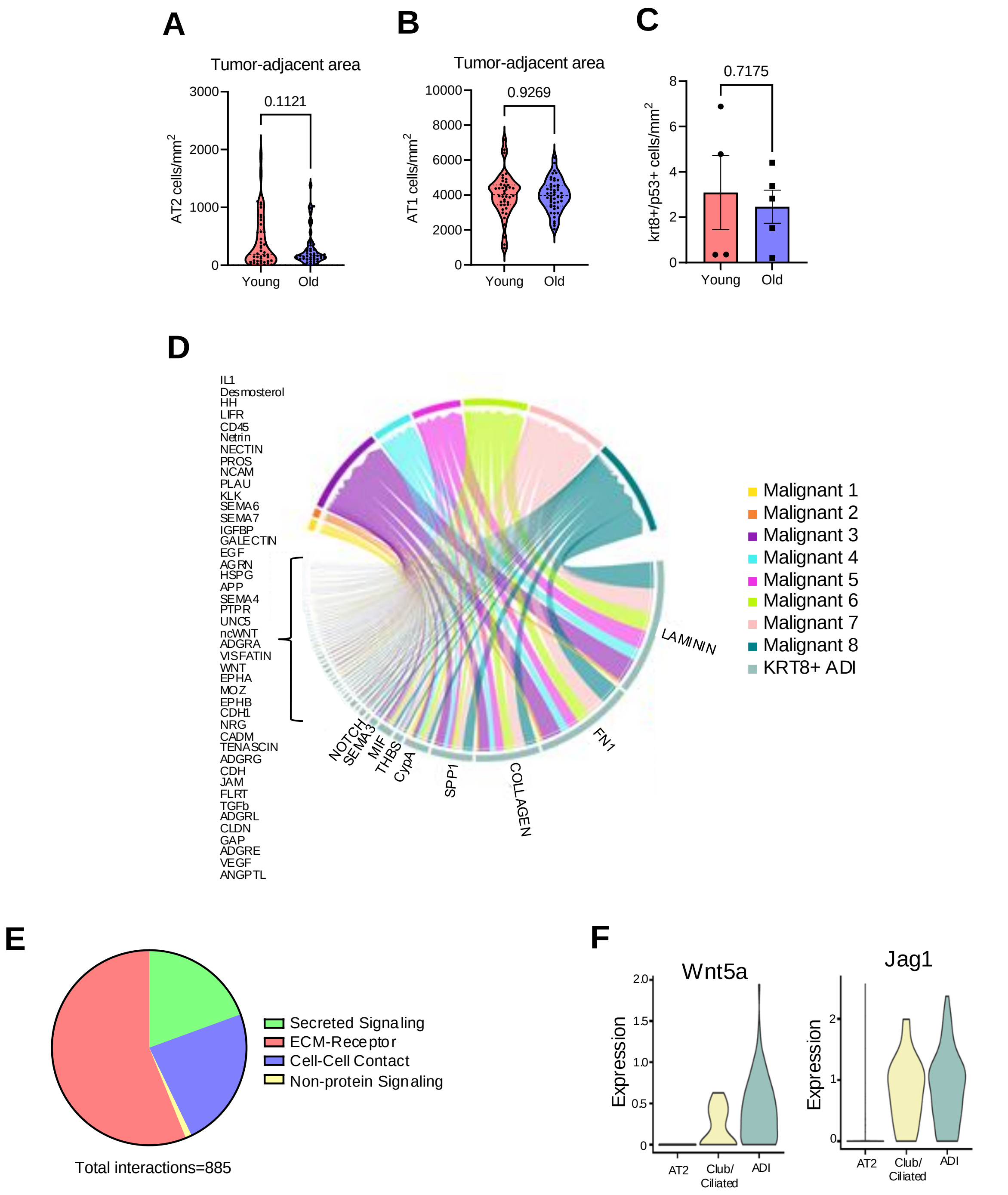
